# Cell-free mtDNA drives pyroptosis and inflammation in m.3243A>G mitochondrial disease models

**DOI:** 10.64898/2026.06.03.729489

**Authors:** Monica Moresco, Alessandro Rapone, Concetta Valentina Tropeano, Mariantonietta Capristo, Giada Capirossi, Danara Ormanbekova, Claudio Fiorini, Alberto Pietro Pasti, Francesco Valle, Alberto Danese, Simone Patergnani, Leonardo Caporali, Chiara La Morgia, Anu Suomalainen, Paolo Pinton, Marco Tigano, Valerio Carelli, Alessandra Maresca

**Affiliations:** IRCCS Istituto delle Scienze Neurologiche di Bologna, Programma di Neurogenetica, Bologna, Italy; Department of Biomedical and Neuromotor Sciences (DIBINEM), University of Bologna, Bologna, Italy; Consiglio Nazionale delle Ricerche - Istituto per lo Studio dei Materiali Nanostrutturati, Bologna, Italy; Department of Medical Sciences, Section of Experimental Medicine and Laboratory of Technologies for Advanced Therapy (LTTA), University of Ferrara, Ferrara, Italy; Research Programs Unit, Stem Cells and Metabolism, University of Helsinki, Helsinki, Finland; HUS Diagnostic Center, Helsinki University Hospital, Helsinki, Finland; HiLife, University of Helsinki, Helsinki, Finland; Maria Cecilia Hospital, GVM Care & Research, Cotignola, Italy; Department of Pathology & Genomic Medicine, Thomas Jefferson University, Philadelphia, Pennsylvania, USA

**Keywords:** cell-free mtDNA, inflammation, pyroptosis, MELAS, mitochondrial dysfunction

## Abstract

Mitochondrial Encephalopathy, Lactic Acidosis, and Stroke-like episodes (MELAS) syndrome is primarily caused by the heteroplasmic m.3243A>G/*MT-TL1* pathogenic variant. Patients exhibit elevated circulating cell-free mtDNA (cf-mtDNA) in plasma, which acts as a damage-associated molecular pattern. Using patient-derived fibroblasts and neuronal progenitors, as well as transmitochondrial cytoplasmic hybrids (cybrids), we show that mutant cells release higher levels of cf-mtDNA than *wild-type* controls, demonstrating that the m.3243A>G pathogenic variant drives mtDNA release. Mechanistically, increased mitochondrial oxidative stress promotes mtDNA oxidation and fragmentation, leading to Ca^2+^ overload and subsequent mtDNA extrusion. This, in turn, triggers inflammasome activation and pyroptosis, resulting in the secretion of pro-inflammatory cytokines and the activation of innate immune pathways. Pharmacological inhibition of the Mitochondrial Calcium Uniporter (MCU) or Voltage-Dependent Anion Channel (VDAC) reduced mtDNA release, confirming their involvement. Overall, our findings reveal a previously unrecognized mechanism in MELAS linking mitochondrial dysfunction to innate immune activation, with potential implications for therapeutic intervention.

## Introduction

Mitochondrial diseases are among the most common inherited metabolic disorders, despite many of them being rare or ultra-rare when taken individually. They can be caused by pathogenic variants in either nuclear or mitochondrial DNA (mtDNA), and characterized by marked clinical and genetic heterogeneity ^1^.

The m.3243A>G/*MT-TL1* nucleotide change, affecting the gene encoding for the tRNA Leucine (UUR) (tRNA^Leu(UUR)^) is the most prevalent heteroplasmic mtDNA pathogenic variant ^2^ and is classically associated with Mitochondrial Encephalomyopathy, Lactic Acidosis, and Stroke-like episodes (MELAS) ^3^, a mitochondrial disease phenotypically defined in 1984 ^4^. However, this variant also underlies a broad spectrum of related phenotypes, depending on variable heteroplasmy levels in different tissues and cells, as well as on unknown genetic modifiers. This spectrum of disorders ranges from milder phenotypes such as maternally inherited diabetes and deafness (MIDD) ^5^, or mitochondrial myopathy with chronic progressive external ophthalmoplegia (CPEO) to more severe presentations including stroke-like episodes (SLEs), and multisystem involvement ^6,7^.

At the molecular level, the m.3243A>G variant impairs tRNA^Leu(UUR)^ function, ultimately affecting mitochondrial protein synthesis ^8–11^, leading to defective oxidative phosphorylation, increased reactive oxygen species (ROS) production, and bioenergetic failure ^12,13^, predominantly involving Complex I ^14^. Although mitochondrial dysfunction and oxidative stress are well-established features of MELAS, the transition from this general pathogenic mechanism to clinical, and tissue-specific features remains poorly understood, including the genesis of SLEs.

In addition to their canonical role in cellular metabolism, mitochondria are increasingly recognized as key regulators of innate immunity ^15^. In particular, the mtDNA, when released into the cytoplasm or extracellular space, can act as a damage-associated molecular pattern (DAMP), engaging innate immune pathways including cGAS–STING, inflammasome signaling, and interferon-mediated responses ^16,17^. Circulating cell-free mtDNA (cf-mtDNA) has been implicated in a range of pathological conditions, contributing to sterile inflammation and disease progression 18.

We and others recently reported elevated levels of circulating cf-mtDNA in patients carrying the m.3243A>G/*MT-TL1* variant, suggesting a potential role for mtDNA release in MELAS pathogenesis ^19–21^. However, the mechanisms driving mtDNA release and its functional consequences in this context remain largely unexplored.

Here, we investigate the origin and impact of cf-mtDNA in different cellular models carrying the m.3243A>G/*MT-TL1*, including patient-derived fibroblasts and neuronal progenitors, as well as transmitochondrial cybrids ^22^. We describe a mechanism linking mitochondrial oxidative stress to mtDNA oxidation, fragmentation, and Ca^2+^-dependent extrusion, ultimately leading to activation of innate immune pathways. These findings uncover a previously unrecognized link between mitochondrial dysfunction and inflammation in MELAS, with potential implications for therapeutic intervention.

## Materials and Methods

### Cell lines and culture conditions

Fibroblast cell lines were established from skin biopsies, after having obtained informed and written consent from patients and controls for the study and for all procedures. MELAS fibroblasts were generated from two male patients (F#1 and F#2) carrying the m.3243A>G variant at 86% and 96% of heteroplasmy, respectively, and compared with age and sex-matched fibroblasts derived from three healthy donors.

MELAS transmitochondrial cytoplasmic hybrids (cybrids, derived from 143B.TK− osteosarcoma cell line, 3243-C) were kindly provided by Eric Schon, whereas the two control cybrid cell lines were generated from healthy individuals. Fibroblasts and cybrids were cultured in Dulbecco Modified Eagle Medium (DMEM, GIBCO) supplemented with 10% fetal bovine serum (FBS, GIBCO), 2 mmol/l L-glutamine, 100 units/ml penicillin and 100 μg/ml streptomycin, at 37°C in a 5% CO2 humidified incubator. The metabolic stress condition medium was represented of DMEM glucose-free supplemented with 5 mmol/L galactose, 5 mmol/L Na-pyruvate, and 5% FBS.

### NPCs generation

Human induced pluripotent stem cells (hiPSCs) derived from a patient carrying 75% of m.3243A>G heteroplasmy previously generated and characterized ^23^, together with their isogenic control, were expanded in StemMACS PSC-Brew XF Medium (130-127-865 Miltenyi Biotec), supplemented with Y27632 10µM (HY-10583-10mg, MedChemExpress) after every passage. hiPSCs were differentiated into NPCs according to Choi et al. ^24^, with minor modification ^25^. To form Embryoid Bodies (EBs), hiPSC were dissociated as single cells using Accutase (A6964-100ML, Sigma-Aldrich), and incubated at 37° in Anti-Adherence (07010, Stem Cell Technologies) coated six-well plates for 3-4 days with Differentiation Media (DM) consisting of 1:1 solution of DMEM F12 (21331020, Thermo-Fisher Scientific) and Neurobasal (21103049, Thermo-Fisher Scientific), supplemented with N2 1X (17502001, Thermo-Fisher Scientific), B27 1X (12587010, Thermo-Fisher Scientific), Glutamax 2 mM (35050061, Thermo-Fisher Scientific), MycoZapTM Plus 1X (LOVZA-2012, Euroclone), EGF 10 ng/ml (HY-P7190, MedChemExpress) bFGF2 10 ng/ml (3718-FB RD-Systems). The EBs were dissociated and seeded on Matrigel (356231, Corning) coated 6-well plates in Expansion Media (EM), consisting of DM without Neurobasal and supplemented with BSA 1,5 mg/ml (A9418-50g Sigma-Aldrich). After one week cells were dissociated with Accutase and expanded for 3-4 passages in order to obtain a homogeneous NPCs culture. NPCs were characterized via immunofluorescence for the expression of SOX2 (1:300, Proteintech, 101064-1-AP) and NESTIN (1:100, Stem Cell Technologies, 60091) markers.

### Cell-free mtDNA assessment

Fibroblasts were seed in a 24-well plate at concentration of 5×10^4^ cell/well in stress condition and of 2×10^4^ cell/well in standard condition. Cybrids were seed in in a 24-well plate at concentration of 3,5×10^4^ cell/well in metabolic stress condition and of 1,5×10^4^ cell/well in standard condition. NPCs were seeded in matrigel-coated 24-well plates at a concentration of 3×10^4^ cells/well in EM + Y27632 10 µM. After 24 h, fibroblasts and NPCs were incubated with standard medium for 48 h, while cybrids were incubated for 16 h. Fibroblasts and cybrids were also incubated in metabolic stress (glucose-free medium supplemented with 5mM galactose) condition for 48 h or 16 h, respectively. At the time points the media were collected, centrifuged at 4 ° C, 10.500 g, for 10 min.

Cell-free DNA (cf-DNA) was extracted from 500 μl of condition medium using the MagMAX Cell-Free DNA Isolation Kit (A29319, Thermo Fisher Scientific), following manufacturer’s instruction. The cf-mtDNA was quantified by droplet digital-PCR (ddPCR, QX200™ Droplet Digital™ PCR System, BIO-RAD) with Taqman-based methods, as previously described ^19^, and expressed as copies of target gene on μl of template analyzed (copies/μl template) and normalized to protein content measured by SRB assay.

### MtDNA content and m.3243A>G variant heteroplasmy assessment

In parallel with cf-mtDNA assessment we seeded in a 24-well plate fibroblast and cybrids at concentration of 5×10^4^ cell/well and of 2×10^4^ cell/well (3,5×10^4^ cell/well in metabolic stress condition), respectively for total DNA extraction. DNA was isolated using a commercial kit following manufacturer’s instructions (Nucleospin Tissue kit for DNA from cells and tissue, Macherey-Nagel). MtDNA content was assessed by real time-PCR, as previously described ^19^. MtDNA and cf-mtDNA heteroplasmy levels were assessed by ddPCR using Prime PCR Custom Assays (Bio-Rad), discriminating the wild-type genomes from the mutated ^19^. Heteroplasmy was expressed as percentage of mutated mtDNA genomes on total mtDNA.

### Assessment of cytoplasmic mtDNA and dsRNA by immunofluorescence

Fibroblasts were seeded at 1.5 × 10 cells/well on coverslips in 4-well plates and, after 48 h, fixed with 4% paraformaldehyde for 20 min.

For mtDNA staining after permeabilization and blocking with PBS containing 5% FBS and 0.25% Triton X-100 for 2 h, mitochondria were labeled by incubating cells with an anti-TOM20 antibody (1:200; Proteintech, 11802-1-AP) and an anti-DNA antibody (1:100; Millipore, CBL186), followed by incubation with Alexa Fluor 488– and Cy3-conjugated secondary antibodies for 1 h. After washing, cells were incubated with Hoechst (1:1000) for 10 min, washed again, and mounted on coverslips. For dsRNA staining, fibroblasts were permeabilized for 5 min at room temperature with a buffer containing 0.5% Triton X-100, 20 mM HEPES, 50 mM NaCl, 3 mM MgCl, and 300 mM sucrose, and then incubated in blocking buffer (0.1% Triton X-100, 1% BSA, and 0.2% gelatin in PBS) for 1 h. Primary antibodies—anti-dsRNA J2 (1:2000; Cell Signaling Technologies, BK76651L) and anti-TOM20 (1:5000; Proteintech, 11802-1-AP)—were diluted in blocking buffer and applied overnight at 4°C, followed by incubation with Cy5-conjugated anti-rabbit and Alexa 488–conjugated anti-mouse secondary antibodies (1:200) for 1 h at room temperature. Images were acquired using a Nikon A1 confocal microscope with a 60× oil-immersion objective, and colocalization analysis was performed on maximum-intensity projections using ROIs to isolate individual cells. Colocalization parameters, including Mander’s overlap coefficient (MOC) and the k and k coefficients, were calculated using NIS-Elements AR software.

### Cell Viability

Cell viability was assessed by colorimetric Sulforhodamine B (SRB) assay, after galactose-medium incubation. Briefly, fibroblasts and cybridswere seeded in 24-well plate at concentration of 5×10^4^ cell/well and of 3,5×10^4^ cell/well, respectively. After 24 h, the cells were incubated in metabolic stress condition. Fibroblast cells were then fixed at 0 h, 24 h, 48 h and 72 h whereas cybrid cells were fixed at 0 h, 8 h, 16 h, and 24 h with 10% trichloroacetic acid (Merck, T6399) for 1 h at 4 °C and stained with 0.4% SRB (Merck, S1402) in 1% acetic acid. After 30 min of incubation, wells were washed with 1% acetic acid and the dye was solubilized with 10 mM Tris pH 10.5. Absorbance at 564 nm was then acquired by Enspire microplate reader instrument (Revvity).

### Mitochondrial respiration evaluation

Oxygen consumption rate (OCR) was measured in adherent fibroblasts (2×10^4^ cells/well density)and NPCs (8x10^4^ cells/well cells/well) with the XFe24 Extracellular Flux Analyzer (Seahorse, Agilent Technologies) following previously described protocol ^26^. OCR was normalized to protein content, as cell number surrogate, measured by SRB assay. Basal, ATP-linked and maximal respirations were calculated as previously described ^26^.

### Evaluation of mitochondrial membrane potential

Mitochondrial membrane potential was evaluated using JC-1 fluorescent probe (Thermo Fischer). Briefly, 8×10^3^ cells and 1,6×10^4^ cells in standard and stress condition respectively were seeded on a 96-well clear bottom black plate. The day after cells were incubated with the standard or stress condition medium for 48 h. At the time point JC-1 at 5 ug/ml concentration was added for 30 minutes at 37 °C. Then, cells were washed with PBS and medium without phenol red in standard and stress condition replaced for fluorescence measurement at 529 nm and 590 nm acquired by Enspire microplate reader instrument (Revvity).

### Transcriptome library preparation and analysis

Total RNA was extracted from fibroblast cell lines using PureLink™ RNA Mini kit (12183018A, Invitrogen) and treated with DNaseI (AMPD1-KT, Sigma-Aldrich). For RNAseq, sample libraries were prepared from 800 ng of RNA input using the TruSeq Stranded mRNA kit (Illumina), following manufacturer’s instructions. Next generation sequencing (NGS) libraries’ sizing was assessed on a 5200 Fragment Analyzer system (Agilent), then sequencing was performed on a NovaSeq 6000 instrument (Illumina) as 2x100 bp paired-end reads.

The fastq files were generated by demultiplexing the libraries using bcl2fastq v2.20. The raw reads were trimmed using Trimmomatic-0.39 ^27^ and the quality of reads was checked using FastQC v0.11.5. The RNAseq libraries were aligned to hg38 (genome_release_39_CRCh38.p13) using the STAR2 with the following parameters; twopassMode Basic – quantModeTranscriptomeSAM ^28^. The replicates were then merged and the read count matrix was generated using salmon -quant v1.5.1 ^29^. Only the genes/transcripts that were supported by at least 10 reads were kept.

The differential expression analysis was performed using *DESeq2* ^30^. Estimated log2 fold changes (log2FC) were shrunk using the Bayesian shrinkage estimator *apeglm* ^31^, and *p*-values associated with fold changes were adjusted for false discovery rate (FDR) using the Benjamini–Hochberg correction. Adjusted *p*-values < 0.05 were considered statistically significant. Normalized count data were obtained via regularized logarithm (*r*log) transformation to remove dependence of variance on the mean and to account for library size. Fast Gene Set Enrichment Analysis was performed using the *fgsea*package (https://bioconductor.org/packages/release/bioc/html/fgsea.html), with gene sets retrieved from the Broad Institute’s MsigDB collection via the *msigdbr* package ^32^. We employed the fgseaMultilevel function and the epsilon (*eps*) parameter was set to a lower limit of 0. To investigate mitochondrial gene deregulation, the differential expression dataset was enriched using Integrated Mitochondrial Protein Index (IMPI – human genes collection, fourth and last version of the database was selected: “IMPI-2021-Q4pre”; ^33^. From IMPI database, we kept the genes encoding verified mitochondrial proteins with “gold standard” evidence of mitochondrial localization, the ones encoding associated mitochondrial proteins with evidence of mitochondrial localization, but lacking visual confirmation, and the genes encoding ancillary mitochondrial proteins with no evidence of mitochondrial localization, but reported to affect mitochondrial function or morphology. We performed the enrichment analysis of the IMPI-enriched dataset using the *enrichR* package ^34^. All statistical analyses and graphical representations were performed using R version 4.5.1 and Bioconductor version 3.21. Finally, gene expression data were analyzed using Ingenuity Pathway Analysis software (Qiagen) for manual inspection of target pathways of interest.

### Live-cell assays for the determination of caspase activity

Briefly, fibroblasts were seeded into clear-bottom white 96-well plates at a density of 8 × 10³ cells/well under standard culture conditions and 12 × 10³ cells/well under metabolic stress conditions. The following day, the culture medium was replaced with phenol red-free standard or metabolic stress medium, and cells were incubated for an additional 48 h before the assays were performed according to the manufacturers’ instructions. Caspase-3/7 activity was measured in live cells using the ApoLive-Glo™ Multiplex Assay (Promega, G6410). Caspase-8 and caspase-9 activities were determined in live fibroblasts using the Caspase-Glo® 8 (Promega, G8201) and Caspase-Glo® 9 (Promega, G8211) assays, respectively, under the same experimental conditions and according to the manufacturers’ protocols. Specific caspase-1 activity was assessed using the Caspase-Glo® 1 Inflammasome Assay (Promega, G9951) according to the manufacturer’s instructions. Fluorescence and luminescence signals were acquired using an EnSpire multimode microplate reader (Revvity). Caspase-8, 9 and 1 activities were normalized to total protein content determined by the SRB assay.

### AlphaLISA assay for cytokines quantification

Fibroblasts and NPCs were seeded under the same conditions as those used for the cf-mtDNA assessment. Then, medium was collected and centrifuged for removal of cell debris and frozen at -80 until analysis. AlphaLISA assays were used to evaluate in the medium INFα (AL297C, Revvity), TNFα (AL3157C, Revvity), IL-6 (AL223C, Revvity), IL-1β (AL3160C, Revvity) and IL-18 (AL3137C, Revvity), following manufacturer’s instructions. The signal at λ_ex/em_ of 680/615 nm was acquired by Enspire microplate reader instrument (Revvity). The data obtained were normalized by SRB assay.

### SDS-PAGE and immunoblotting

Total lysates were prepared using RIPA lysis buffer and a standard protocol ^35^. Proteins were separated on pre-cast NuPAGE 4–12% bis-tris glicyne gels (Life Technologies) and then transferred on nitrocellulose membranes, using the XcellSure Lock (Life Technologies) apparatus. After blocking with 5% milk, membranes were blotted with primary antibodies specific for: GAPDH (Santa Cruz Biotechnologies sc-47724, dilution 1:5000), ITPR3 (Abclonal A23202, dilution 1:3000), MCU (Proteintech 26312-1-AP, dilution 1:3000), RIG-I (D14G6, Cell Signaling #3743, dilution 1:1000), phospho-STAT1 (58D6, Cell Signaling #9167, dilution 1:700), TOM20 (Proteintech, 11802-1-AP, dilution 1:5000), TUBB (Abnova, H00203068-M04, dilution 1:10000), Catalase (AB Clonal, A11777, dilution 1:500), GPX1 (Proteintech 29329-1-AP, dilution 1:500), MnSOD (Merck Millipore #06-984, dilution 1:600), STING (Proteintech 66680-1-Ig, dilution 1:5000), MAVS (Cell Signaling, #E993, dilution 1:1000), MGME1 (Proteintech, 23178-1-AP, dilution 1:1000), FEN1 (Santa Cruz, SC-28355, dilution 1:500).

Fluorescent secondary antibodies anti-rabbit or anti-mouse (Licor, 1:7000) were used for immunodetection using the Licor Odyssey instrument (LICORbio).

### Assessment of mitochondrial Ca^2+^

To assess resting mitochondrial Ca^2+^ levels, fibroblasts and cybrids were transfected with a plasmid encoding mtGCaMP6m. After 36 h, coverslips were imaged using an Olympus Cell^R high-resolution multi-wavelength fluorescence microscope equipped with a 40× oil-immersion objective. Basal mtGCaMP change of fluorescence (494/406) was recorded for 1 min under each experimental condition. The same experimental procedure was used to measure mitochondrial Ca^2+^ levels following stimulation with a Ca^2+^-mobilizing agonist. In these experiments, cells were challenged with 100 µM ATP to induce Ca^2+^ release from the endoplasmic reticulum, and the resulting changes in mitochondrial Ca^2+^ were monitored by fluorescence microscopy.

### Flow cytometry measurement of reactive oxygen species production

Intracellular and mitochondrial reactive oxygen species (ROS) were quantified by flow cytometry using an Attune™ NxT Acoustic Focusing Cytometer (Thermo Fisher Scientific). Mitochondrial superoxide production was assessed with MitoSOX™ Red (Thermo Fisher Scientific, M36008), while total cellular ROS levels (hydrogen peroxide, peroxynitrite and derived oxidizing radicals) were measured using 2’,7’-dichlorodihydrofluorescein diacetate (H DCF-DA, Sigma Aldrich, D6883). Briefly, cells were harvested, washed in pre-warmed PBS, and incubated with 2.5 µM MitoSOX Red or 2 µM H DCF-DA for 30 min at 37 °C in the dark, according to the manufacturer’s instructions. Following incubation, cells were immediately acquired on the Attune™ cytometer using standard excitation (Ex)/emission settings (Em) (MitoSOX™ Red: Ex 510 nm/Em 580 nm; H DCF-DA: Ex 488 nm/Em 530 nm). A minimum of 5,000 events/gated per sample was collected, excluding debris and doublets based on forward and side scatter parameters. Data were analyzed using Attune™ Cytometric Software, and mean fluorescence intensity (MFI) normalized on cell count was used as a quantitative measure of ROS levels.

### Cell fractioning

Cell fractionation was performed in cells line, as previously described ^36^. Briefly, the cells pellet was suspended in homogenization buffer (200 mM mannitol, 70 mM sucrose, 1 mM ethylene glycol-bis (β-aminoethylether)-N,N,N,N-tetraacetic acid (EGTA), 4-(2-hydroxyethyl)-1-piperazineethanesulfonic acid (HEPES), pH 7.4) and gently disrupted by automated Dounce homogenization. The homogenate was centrifuged to remove nuclei and unbroken cells, and the resultant supernatant was centrifuged (12,000g for 15 minutes at 4 °C) to pellet crude mitochondria and cytosolic fraction (supernatant). To collect pure mitochondria, crude mitochondria were resuspend in buffer and centrifuged again at 12,000g for 15 minutes at 4 °C. After centrifugation, the mitochondrial fraction was freeze at -80 °C for further analysis (e.g. DNA extraction).

### Mitochondrial DNA damage assessment

MtDNA was extracted from pure mitochondria using the Macherey-Nagel NucleoSpin™ Tissue kit (cat. Number 740952.50), following the manufacturer’s instructions.

The EpiQuik™ 8-OHdG DNA Damage Quantification Direct Kit (Colorimetric, P-6003-96, EpigenTek, USA) was used to quantify 8-hydroxy-2’-deoxyguanosine (8-OHdG), an oxidized derivative of deoxyguanosine generated by hydroxyl radicals, singlet oxygen, and one-electron oxidants in cellular DNA

MtDNA fragmentation was assessed by capillary electrophoresis on a 5200 Fragment Analyzer System using the Large Fragment Kit (Agilent). Sizing of the DNA was confirmed as being compatible with mtDNA only (<17 kb), then we calculated the ratio of peak areas for fragments smaller or larger than 2 kb.

For Atomic Force Microscopy (AFM), a 10 μl aliquot containing mtDNA (typically a 10-fold dilution of the stock solution) and 2 mM MgCl_2_ final concentration was deposited on a freshly cleaved mica surface. The sample was then incubated 10 min, letting the DNA adsorb onto the substrate ^37^. Afterward, the surface was gently rinsed with 1 ml Milli-Q water (18.2 MΩ.cm resistivity) and then blow-dried with Nitrogen.

Images were collected using a MultiMode VIII SPM with a Nanoscope V controller (Bruker Instruments, Santa Barbara, CA, USA) operated in ScanAsyst mode in air. In this mode of operation several force distance curves are performed on each sample point controlling the maximum applied load. The image collected is the profile measured by the scanner to keep the force constant. The AFM cantilevers used had a nominal spring constant of 0.2 N·m^−1^ (Bruker ScanAsyst air) and tip radius of 2 nm. All the recorded AFM images consist of 512 × 512 pixels with scan frequency ≤1 Hz. Images analysis was performed using the Gwyddion software (Version 2.25); briefly: each image was flattened to remove the substrate slope and then the DNA molecule length of each molecule was measured using the segmentation tool.

### Pharmacological inhibition of MCU and VDAC oligomerization

Cybrids were seeded in 24-well plates at a density of 1.5×10 cells/well under standard culture conditions. The following day, cells were treated with the MCU inhibitor MCU-i11 (Cat. No. HY-W194810) at 2 μM or with VBIT-4 (Cat. No. HY-129122) at 10 μM for 16 h. After treatment, the culture medium was collected, centrifuged, and stored at −80 °C until cf-mtDNA analysis.

### Statistical Analysis

GraphPad Prism for Windows (GraphPad Software) was used for statistical analyses. Comparisons of control and MELAS mutant were assessed by paired or unpaired t tests with Welch’s or FDR correction. Differences between controls and MELAS mutant cell lines were analyzed using one-way or two-way ANOVA and Dunnett’s or Šidák multiple comparisons tests. For all analyses, differences were considered significant at a P value ≤ 0.05.

## Results

### The release of mutant cf-mtDNA characterizes m.3243A>G fibroblasts

Starting from our seminal finding that patients carrying the m.3243A>G/*MT-TL1* pathogenic variant (herein m.3243A>G), associated with different clinical phenotypes (MELAS and prominent muscular phenotype without SLEs) present increased levels of ccf-mtDNA in plasma ^19^, we first evaluated mtDNA release in patient-derived fibroblasts.

To this aim, we used two fibroblast lines derived from patients with discordant phenotypes: one patient never experienced SLEs (F#1, bulk heteroplasmy of 85.3% ± 2.02 standard deviation [SD]), whereas the other patient had a frequent recurrence of SLEs (F#2, bulk heteroplasmy of 94.0% ± 2.36 SD). For details on clinical history and patients’ findings, see supplementary materials (Supplementary Figures S1 A and S2 A-B).

In fibroblasts, we quantified cf-mtDNA released under standard (glucose-containing medium) and stress (glucose-free, galactose-containing medium) conditions after 48 hours of growth. Using galactose as carbon source instead of glucose forces cells to use oxidative phosphorylation (OXPHOS) rather than glycolysis and this is well-known stress condition in the presence of mitochondrial dysfunction ^38^. We observed higher cf-mtDNA levels in both conditions (Fig. 1 A) compared to control cell lines. Furthermore, we compared the m.3243A>G heteroplasmic loads of cf-mtDNA and intracellular mtDNA, showing a preferential release of mutant mtDNA. This latter finding was significant for the F#1 cell line, both in standard and stress conditions, whereas in cell line F#2, we observed significant release of mutant mtDNA compared to intracellular mtDNA exclusively in the stress condition (Fig. 1 B).

**Fig. 1.**
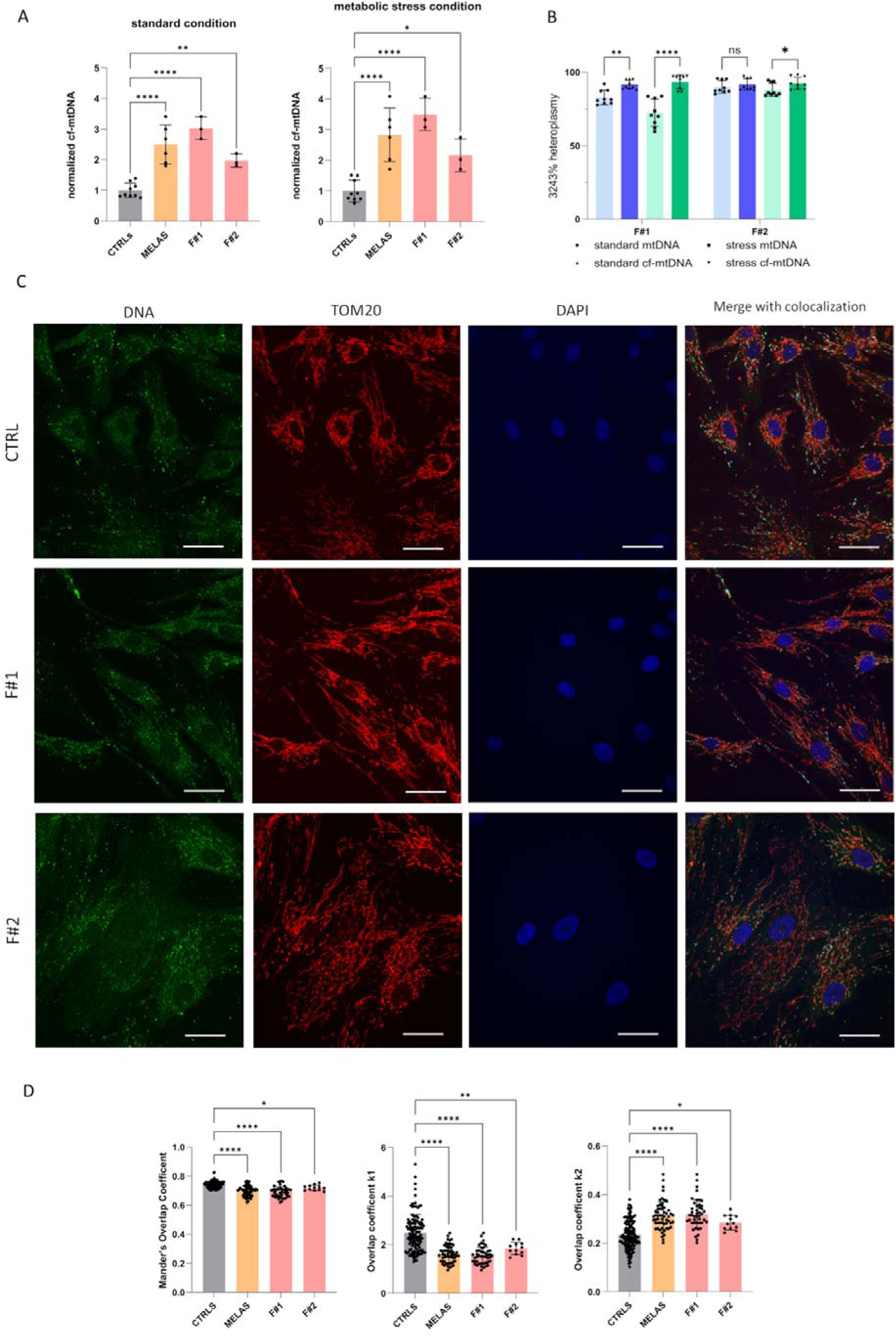
Cf-mtDNA level, heteroplasmy, and co-localization analysis in m.3243A>G fibroblasts. **(A)** The plots show the cf-mtDNA levels in MELAS m.3243A>G-carrying fibroblasts (F#1 and F#2) and matched controls (CTRLS) across 3 independent biological replicates after 48h of growth under standard and metabolic stress conditions. The data were normalized to cell protein content and control means. Data were analyzed by one-way ANOVA with Dunnett’s correction for multiple comparisons; *, **, and **** indicate significant differences from the corresponding control (p-value of 0.02,C0.006, and <0,0001, respectively). **(B)** Heteroplasmy level of m.3243A>G (%) in (on mtDNA) and out (on cf-mtDNA) mutated fibroblasts, in both considered conditions (standard condition in blue and metabolic stress condition in green) for 9 biological replicates. Data were analyzed by a two-way ANOVA test corrected for multiple comparisons (Šidák) and multiple unpaired t tests corrected with false discovery rate (FDR) when appropriate. *, **, and **** indicate significant differences with p-value of 0.05, 0.003, and <0,0001, respectively. **(C)** Representative confocal images showing co-localization in fibroblasts under standard conditions, stained with an anti-DNA antibody (green), anti-TOM20 antibody (red) to label mitochondria, and DAPI (blue) to stain nuclei. Scale bar: 40 μm. **(D)** Overlap coefficient according to Manders (MOC) and overlap coefficients k1 and k2 between MELAS fibroblasts and matched controls following 48h of growth in standard condition. Data were analyzed by one-way ANOVA test corrected for multiple comparisons (Dunnett); *, **, and **** indicate significant differences from the corresponding controls, p-value of 0.04, 0.001 and <0,0001, respectively.

Next, to assess the presence of mitochondria-free mtDNA in the cytosol, we performed immunofluorescence analysis to evaluate the co-localization between mtDNA and a mitochondrial marker (TOM20), as represented by Mander’s Overlap Coefficient (MOC) and the overlap coefficients k1 and k2. We observed a significant loss of co-localization in mutant cells compared to the wild type, as documented by decrease in MOC and the overlap coefficients k1, and an increase in the overlap coefficient k2, indicating that a fraction of mtDNA is in the cytoplasm, thus released from mitochondria (Fig. 1 C and 1 D).

### Mitochondrial dysfunction is a prominent feature of m.3243A>G fibroblasts

To correlate the mtDNA release with mitochondrial dysfunction, we investigated the metabolic profile of the mutant fibroblasts. First, we evaluated cell viability under stress conditions, using glucose-free galactose-containing medium. Both mutant fibroblast lines showed reduced viability after 48 h, with further decline observed after 72 h of growth under these conditions (Fig. 2 A). Next, we investigated the oxygen consumption rate (OCR) in live cells, indicative of oxidative phosphorylation (OXPHOS) functionality (Fig. 2 B). We observed a significant reduction in basal, ATP-linked, and maximum respirations, confirming the metabolic defects in these cell lines, as expected based on previous reports ^10,11,39^. We then assessed mtDNA copy number in the fibroblast cell lines (Fig. 2 C). Both mutant cell lines exhibited an increase of mtDNA content in both standard and stress conditions.

**Fig. 2.**
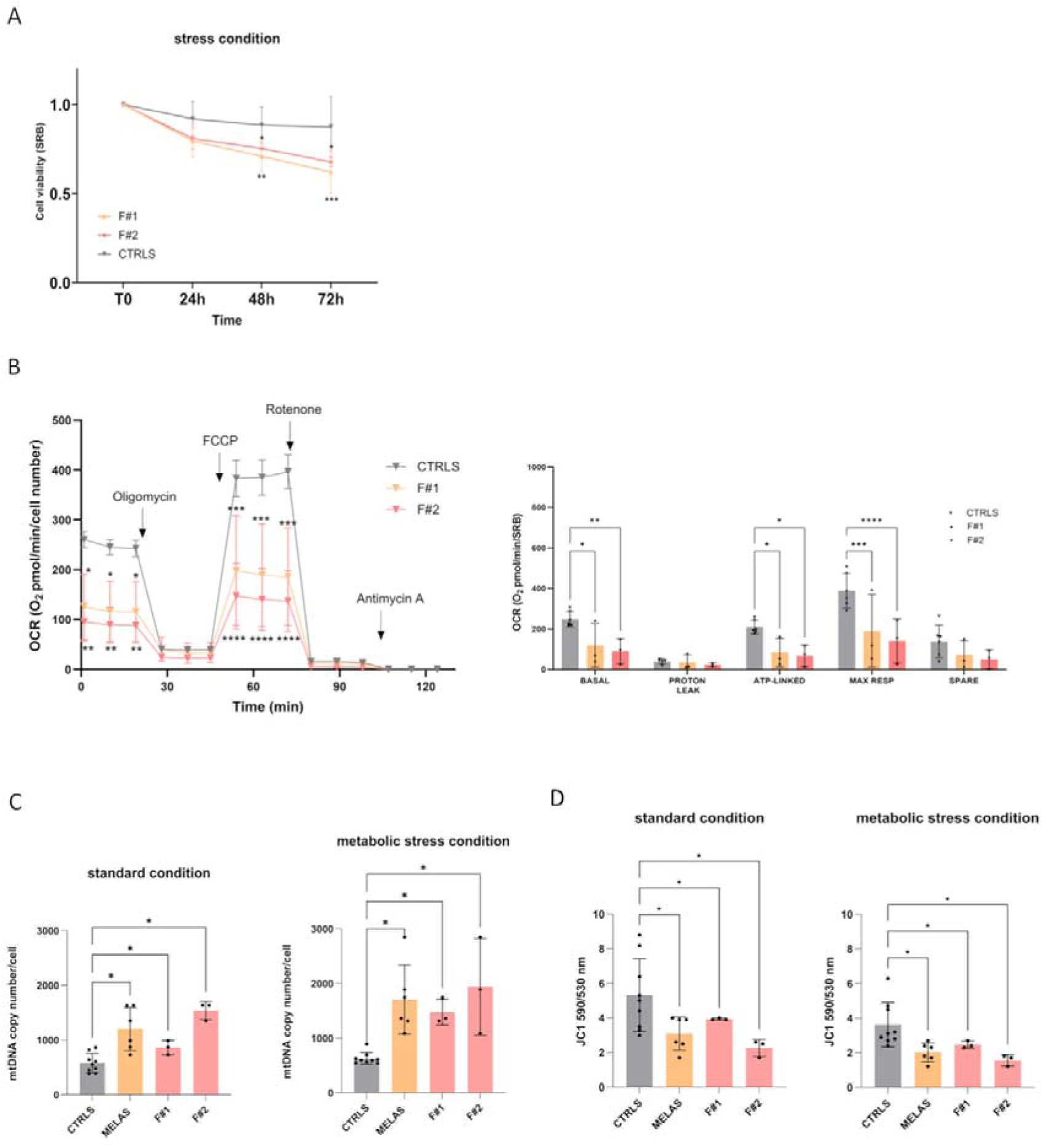
Functional and metabolic characterization of MELAS fibroblasts (F#1 and F#2).(A) Cell viability was assessed by culturing cells under metabolic stress conditions (in a medium containing galactose as the carbon source) for 24, 48, and 72 h. Data are presented as mean ± SD from three independent biological replicates. Statistical analyses were performed using ANOVA followed by Holm-Šídák’s multiple comparisons test. *, ** and, *** values were significantly different from the control cells, pC<C0.03, pC<C0.005, pC<C0.0004, respectively. **(B)** Basal, Proton Leak, ATP-linked, maximal and spare respiration in mutant fibroblasts versus controls. All values are means and SD of 3 independent experiments (biological replicates). Besides, the graphical representation of mitochondrial respiration, OCR was expressed as picomoles O2/min, normalized for protein content, under basal conditions and after injection of OXPHOS modulators (oligomycin, carbonyl cyanide 4-(trifluoromethoxy) phenylhydrazone (FCCP), rotenone, and antimycin A). Data are expressed as meansC±CSD of a minimum of 3 independent experiments (biological replicates). A two-way ANOVA with Dunnett’s correction was used to analyse the data. *,**, ***, ****, corresponding to an adjusted p-value of <0.03, 0.006, 0.0007 and <0.0001, respectively. **(C)** MtDNA copy number was analyzed in MELAS fibroblasts compared to control cells across three independent experiments (biological replicates) under both standard and stress conditions after 48 h of growth. Data were analyzed using the Brown-Forsythe and Welch ANOVA, followed by the Benjamini, Krieger, and Yekutieli multiple-comparison test. * indicates statistically significant discoveries (q < Q). **(D)** The presented plots compare the mitochondrial membrane potential between MELAS fibroblasts and matched controls across three independent biological replicates after 48 h of growth under standard and metabolic stress conditions. Statistical analysis was performed using Brown-Forsythe and Welch ANOVA, followed by the Benjamini, Krieger, and Yekutieli multiple-comparison test. * indicates statistically significant discoveries (q < Q).

Last, we assessed changes in mitochondrial membrane potential using the membrane-permeant dye JC-1. Our results revealed a statistically significant decline in mitochondrial membrane potential in both mutant lines (Fig. 2 D) ^40,41^. In summary, our findings confirm decreased cell viability under galactose medium, mitochondrial respiratory chain deficiency, activation of mitochondrial biogenesis, and reduced mitochondrial membrane potential.

### Transcriptomics highlights pathways related to mtDNA escape, pointing to activation of mitochondrial Ca^2+^ import, oxidative stress, and pyroptosis

To potentially identify molecular pathways upstream and downstream of cf-mtDNA extracellular release, we conducted bulk transcriptomic analysis of the m.3243A>G fibroblast lines under standard conditions. Gene set enrichment analysis of differentially expressed genes (DEGs) in the F#1 and F#2 lines compared with controls revealed significant overrepresentation of cell death pathways—including pyroptosis—as well as inflammation and cytokine secretion. (Fig. 3 A-B). Among the DEGs, we identified several genes of particular interest, including *Interferon Alpha Inducible Protein 27* (*IFI27*), a mitochondrial membrane–associated protein implicated in pyroptosis ^42^. In addition, we observed increased expression of *NLR Family Pyrin Domain Containing 10* (*NLRP10*), a protein that is part of the inflammasome complex and activates pro-inflammatory caspases and induces pyroptosis ^43,44^.

**Fig. 3.**
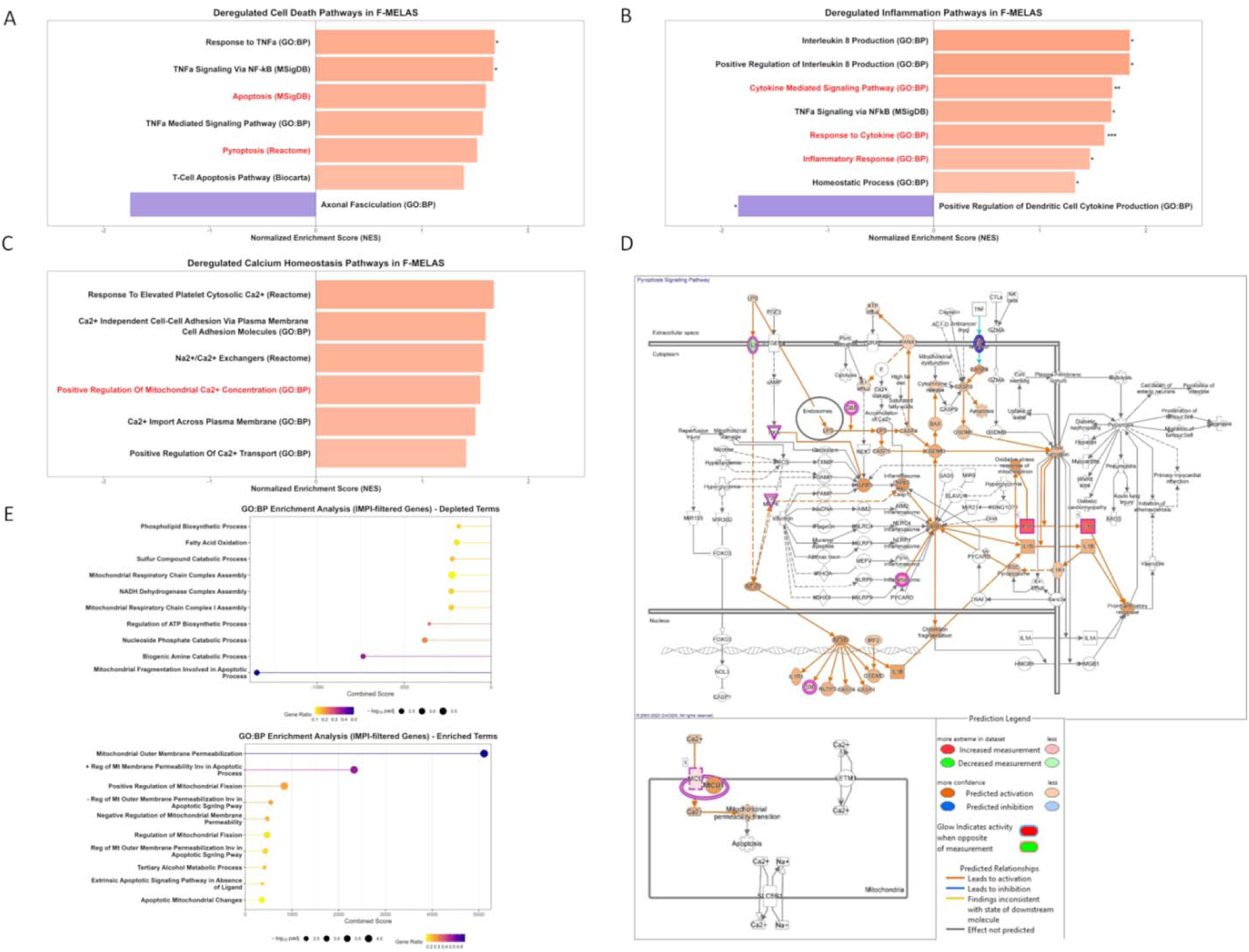
Pathway enrichment analysis of DEGs, predicted activated pathways (IPA), and enrichment of mitochondrial processes. Pathways related to cell death **(A)**, inflammation **(B)**, and dysregulation of Ca^2+^ homeostasis **(C)** are shown. Upregulated processes are highlighted in pink, and down-regulated processes in purple, with corresponding significance levels. **(D)** DEG analysis using IPA revealed a predicted activation of pyroptosis and mitochondrial Ca^2+^ uptake. **(E)** Enrichment analysis was performed using a dataset of mitochondrial genes or genes with mitochondrial-related functions (IMPI).

Moreover, we detected the up-regulation of several pro-inflammatory genes, including *IL-6*, *IL-18*, *IL-32*, and *IL-34*, along with members of the *oligoadenylate synthase* (*OAS*) family (*OAS1*, *OAS2*, and *OAS3*), as well as Toll-like receptors (*TLR*) 1, 5, and 6, and members of the *TNF Superfamily* (*TNFSF*), specifically *TNFSF4*, *TNFSF6*, *TNFSF9*, and *TNFSF15* ^45^. We also observed downregulation of *Suppressor of Cytokine Signaling 2* (*SOCS2*), a classic negative regulator of cytokine signaling that has been described as an anti-inflammatory mediator ^46^.

Furthermore, we observed a reduction in antioxidant activity in MELAS fibroblasts, documented by the downregulation of ROS-detoxifying enzymes such as *catalase* (*CAT*), *microsomal glutathione S-transferase 1 (MGST1)*, and *TBC/LysM domain–containing protein 2 (TLDC2),* which protect both cells and mitochondria from the toxic effects of hydrogen peroxide and lipid hydroperoxides.

In addition, MELAS fibroblasts revealed a consistent upregulation of key genes involved in cellular Ca^2+^ signaling and transport, including *inositol 1,4,5-trisphosphate receptor type 3 (ITPR3)*, a channel that mediates intracellular Ca^2+^ release; *EF-hand domain family member D1 (EFHD1)*; and the *mitochondrial calcium uniporter (MCU)* (Fig. 3 C). Canonical pathway analysis using Ingenuity Pathway Analysis (IPA) confirmed a likely activation of pyroptosis and an increased influx of Ca^2+^ into mitochondria, both pathways associated with the extrusion of mtDNA from mitochondria and its release into the extracellular space (Fig. 3 D).

Finally, gene set enrichment analysis performed on mitochondrial and mitochondrial associated (IMPI dataset, ^33^)- DEGs -highlighted an upregulation of mechanisms such as outer mitochondrial membrane (OMM) permeabilization and associated apoptosis, whereas numerous metabolic processes, including phospholipid biosynthesis, fatty acid β-oxidation, and, as expected, components of the mitochondrial respiratory chain and ATP production, were downregulated (Fig. 3 E).

### Inflammation and sterile immune activation are predominant in MELAS fibroblasts

Transcriptome analysis of MELAS fibroblasts compared with controls revealed potential alterations in innate immunity and inflammation pathways, with a notable predicted activation of pyroptosis, a form of cell death triggered by mtDNA release. To validate these findings, we investigated Caspase-1 activation and the release of pro-inflammatory cytokines into the culture medium. Furthermore, given the apparent interferon-mediated response, we assessed IFNα, TNFα, and IL-6 in the culture medium and the activation of the associated JAK/STAT1 signaling pathway ^47^.

We observed a significant increase in Caspase-1 specific activity in both the F#1 and F#2 cell lines (Fig. 4 A). Downstream of NLRP3 and Caspase-1 activation, the pro-inflammatory cytokines IL-1β and IL-18 are processed and released. In MELAS fibroblasts, we detected elevated levels of these cytokines in the culture medium, although the increase reached statistical significance only in the F#2 cell line (Fig. 4 B-C), suggesting pyroptosis activation. We also observed an upregulation of cell death pathways in the F#2 cell line, accompanied by increased activity of all involved caspases (3/7, 8, and 9), as shown in Supplementary Figure S3 A-C. Similarly, the levels of IFNα, TNFα, and IL-6 were also elevated in MELAS fibroblasts, particularly in F#2 (Fig. 4 D-F), indicating a concurrent activation of two distinct pro-inflammatory signaling pathways. Specifically, this group of cytokines can be released following activation of the cGAS/STING pathway upon recognition of cytosolic mtDNA ^48^, or upon activation of RIG-I/MDA5 and MAVS, sensors of dsRNA ^47^. We failed to detect an increase in STING protein levels (Supplementary Fig. S4A), and, we were unable to assess its possible C-terminal phosphorylation, an event required for the subsequent type I IFN–mediated response ^49^. Regarding the RIG-I/MAVS pathway, we confirmed significantly higher protein levels of these two cellular sensors in both MELAS fibroblast lines (Fig. 4 G–H). Moreover, since this pathway leads to IFNα-mediated STAT1 activation and subsequent induction of interferon-stimulated genes (ISGs) ^42^, we assessed STAT1 phosphorylation and activation by immunoblotting, revealing significantly higher levels in MELAS fibroblasts (Fig. 4 I). We then performed immunofluorescence analysis to evaluate the presence of mitochondrial dsRNA in the cytoplasm. In the F#2 cell line, the co-localization between dsRNA and mitochondria was lower compared to controls, as indicated by the decrease in MOC (shown in supplementary figure S5 A-B), consistently suggesting that a significant fraction of dsRNA is also released from mitochondria and localizes in the cytoplasm. In contrast, the co-localization in the F#1 cell line showed unchanged MOC, but increased k2 (supplementary figure S5 B).

**Fig. 4.**
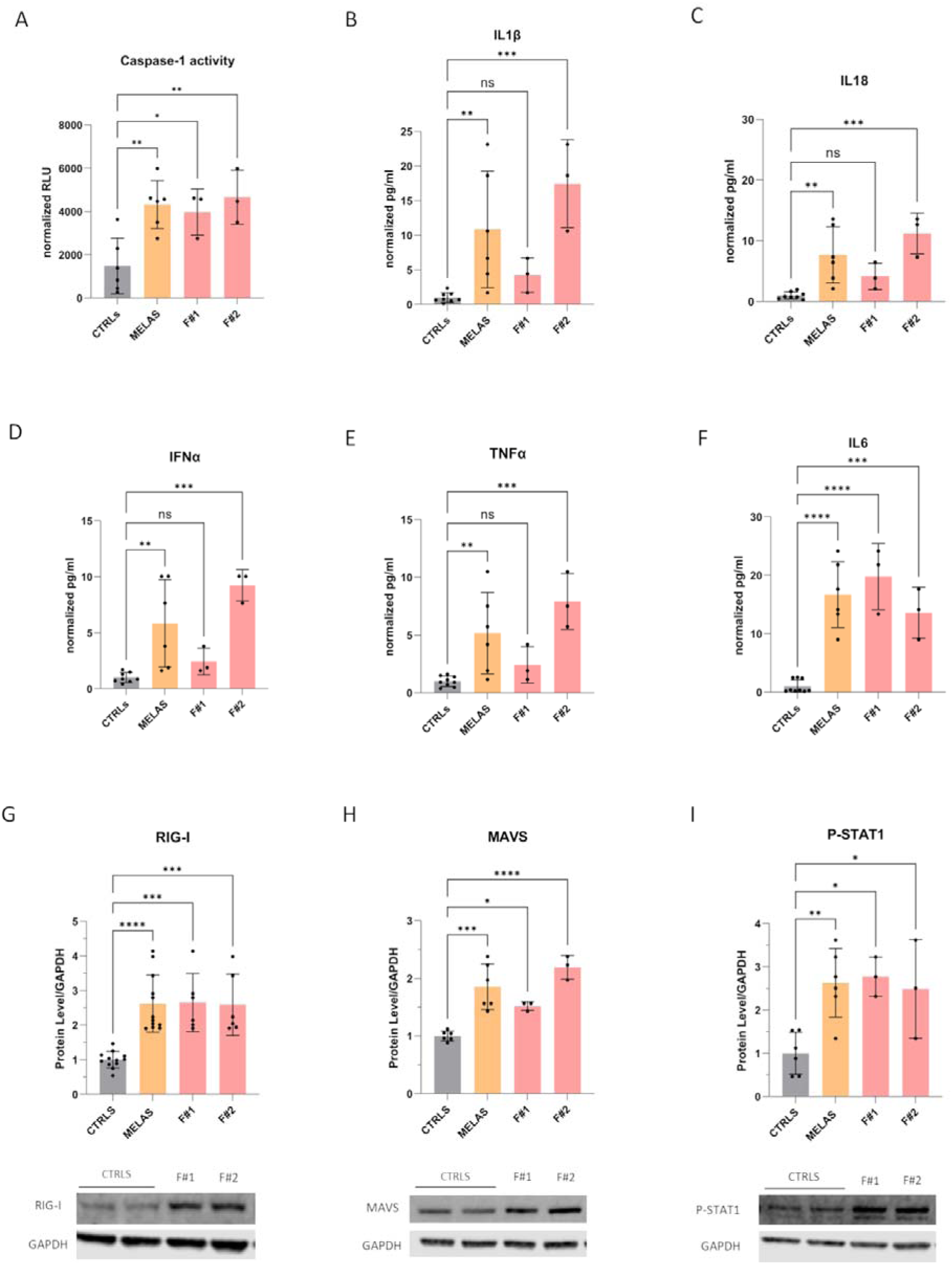
Evaluation of inflammation pathways activated in MELAS fibroblasts.(A) Caspase-1–specific activity was assessed using a commercial luminescence-based kit. The results shown in the graph represent the mean ± SD of three biological replicates *per* cell line, normalized to protein content. One-way ANOVA followed by Dunnett’s multiple-comparison test was used. **(B-F)** Levels of IL-1β, IL-18, IFNα, TNFα, and IL-6 were measured in the fibroblast culture medium after 48 hours of growth under standard conditions, using AlphaLISA assays. The results shown in the graph represent the mean ± SD of three biological replicates *per* cell line, normalized to the mean of the controls. One-way ANOVA followed by Dunnett’s multiple-comparison test was used. **(G–I)** Immunoblot analysis for the evaluation of RIG-I, MAVS, and P-STAT1 protein levels. Representative images and densitometric quantification from at least three independent biological replicates are shown. GAPDH levels were used as a loading control for normalization. For all data presented in this figure, statistical analysis was performed using one-way ANOVA followed by Dunnett’s multiple-comparison test. The symbols *, **, ***, and **** indicate p-values of <0.05, <0.01, <0.001, and <0.0001, respectively.

Overall, these findings demonstrate that the release of mtDNA and mitochondrial dsRNA in MELAS fibroblasts triggers two distinct inflammatory pathways. On one hand, it activates the NLRP3 inflammasome, leading to pyroptosis and the extracellular secretion of the pro-inflammatory cytokines IL-1β and IL-18. In parallel, activation of the RIG-I/MAVS pathway elicits an IFNα response and promotes the release of IL-6 and TNFα, likely through downstream activation of NF-κB and STAT1.

### Enhanced mitochondrial Ca^2+^ uptake and elevated oxidative stress are potential mechanisms triggering the release of mtDNA into the cytoplasm

In addition to the immune and inflammatory responses, transcriptomic analysis also suggested a potential increase in mitochondrial Ca^2+^ levels, along with a less effective defense against oxidative stress. Previous studies have shown that mtDNA oxidation promotes mitochondrial Ca^2+^ uptake, triggering the opening of the mitochondrial permeability transition pore (mPTP) and the subsequent release of oxidized mtDNA ^50,51^. Thus, we analyzed the level of mitochondrial Ca^2+^, and those of the channels mediating its entry: the inner mitochondrial membrane (IMM) channel MCU, and the Mitochondria-Associated Membranes (MAMs) resident receptor ITPR3 ^52^. We detected a significant increase in the levels of both MCU and ITPR3 in MELAS fibroblasts compared to controls (Fig. 5 A-B). Consistently, mitochondrial Ca^2+^ levels, assessed by live-cell imaging with the mtGCaMP sensor ^53^, were higher in MELAS fibroblasts compared with controls (Fig. 5 C). In parallel, we also evaluated the levels of the major enzyme responsible for detoxifying oxygen radicals. The amount of mitochondrial superoxide dismutase (MnSOD) was comparable to, or slightly higher than, those observed in control cells (Fig. 5 D). In contrast, mitochondrial glutathione peroxidase (GPX1) and catalase (CAT), both key hydrogen peroxide (H O)-detoxifying enzymes localized respectively in mitochondria and peroxisomes, were significantly reduced in MELAS fibroblasts relative to controls (Fig. 5 E-F), suggesting a potential accumulation of H O . We next evaluated the general oxidative stress measured by (2’,7’-dichlorodihydrofluorescein diacetate (H DCF-DA) fluorescence, and observed a significantly higher levels of oxidative stress in MELAS fibroblasts compared with controls (Fig. 5 G). In addition, mitochondrial superoxide quantification by mitoSOX™ revealed elevated levels of this radical in the F#2 cell line (Fig. 5 H).

**Fig. 5.**
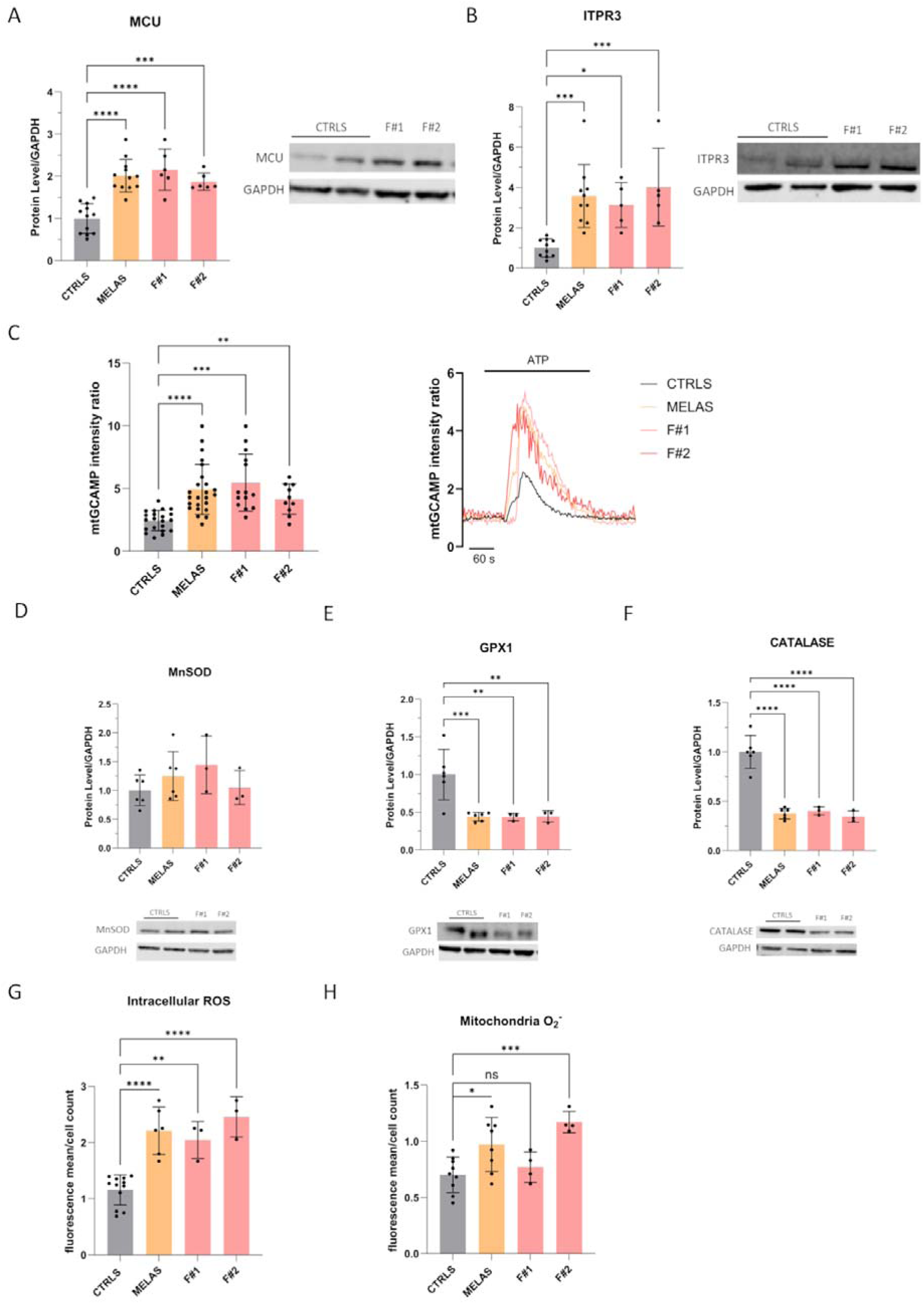
Assessment of mitochondrial Ca^2+^, antioxidant enzymes and reactive oxygen species (ROS).(A–B) Immunoblot analysis of MCU and ITPR3 protein levels. Representative images and densitometric quantification from at least six independent biological replicates are shown. GAPDH levels were used for loading normalization. **(C)** Ca^2+^ uptake was assessed by fluorescence microscopy following transfection with mtGCaMP. Cells were stimulated with ATP to induce Ca^2+^ release from the endoplasmic reticulum. **(D-F)** Immunoblot analysis of antioxidant enzyme levels. Representative images and densitometric quantification from at least three independent biological replicates are shown. GAPDH levels were used for loading normalization. **(G)** Quantification of intracellular ROS levels by flow cytometry, based on 2’,7’-dichlorodihydrofluorescein diacetate (HCDCF-DA) mean fluorescence intensity normalized to cell count, from at least three independent biological replicates. **(H)** Quantification of mitochondrial superoxide by flow cytometry, based on MitoSOX™ mean fluorescence intensity normalized to cell count, from at least four independent biological replicates. Statistical analysis for all data in this figure was performed using one-way ANOVA followed by Dunnett’s multiple comparison test. Symbols *, **, ***, and **** indicate p values of <0.05, <0.01, <0.001, and <0.0001, respectively.

Overall, we validated the alterations identified by the transcriptomic analysis by observing increased mitochondrial Ca^2+^ levels and heightened oxidative stress in mutant fibroblasts.

### Mitochondrial dysfunction drives increased cf-mtDNA release in MELAS neuronal progenitor cells

To confirm our findings in a cellular model more closely resembling the neuronal context, we studied neuronal progenitor cells (NPCs) that we differentiated from previously characterized induced Pluripotent Stem Cells harboring the m.3243A>G pathogenic variant ^23^. Specifically, we analyzed an NPC line with a 75% bulk heteroplasmy level for the m.3243A>G variant and compared it with its isogenic *wild type* control. Both the lines showed expression of the NPCs specific markers SOX2 and Nestin (Supplementary Figure S6).

First, we confirmed the pathogenic phenotype of MELAS NPCs by measuring the OCR in live cells, an indicator of OXPHOS function (Fig. 6 A). MELAS NPCs showed a significant reduction in both basal and ATP-linked respiration, confirming the presence of metabolic defects even in this cell model. In addition, MELAS NPCs exhibited increased mitochondrial superoxide levels compared with control cells (Fig. 6 B). Consistent with these observations, we detected an increased release of mtDNA into the culture medium of MELAS NPCs compared with the isogenic control line (Fig. 6 C). MELAS NPCs exhibited elevated IL-18, IL-6, and TNFα levels compared to their isogenic control (Fig. 6 D-H). IL-1β was below the assay’s detection limit and is therefore not shown.

**Fig. 6.**
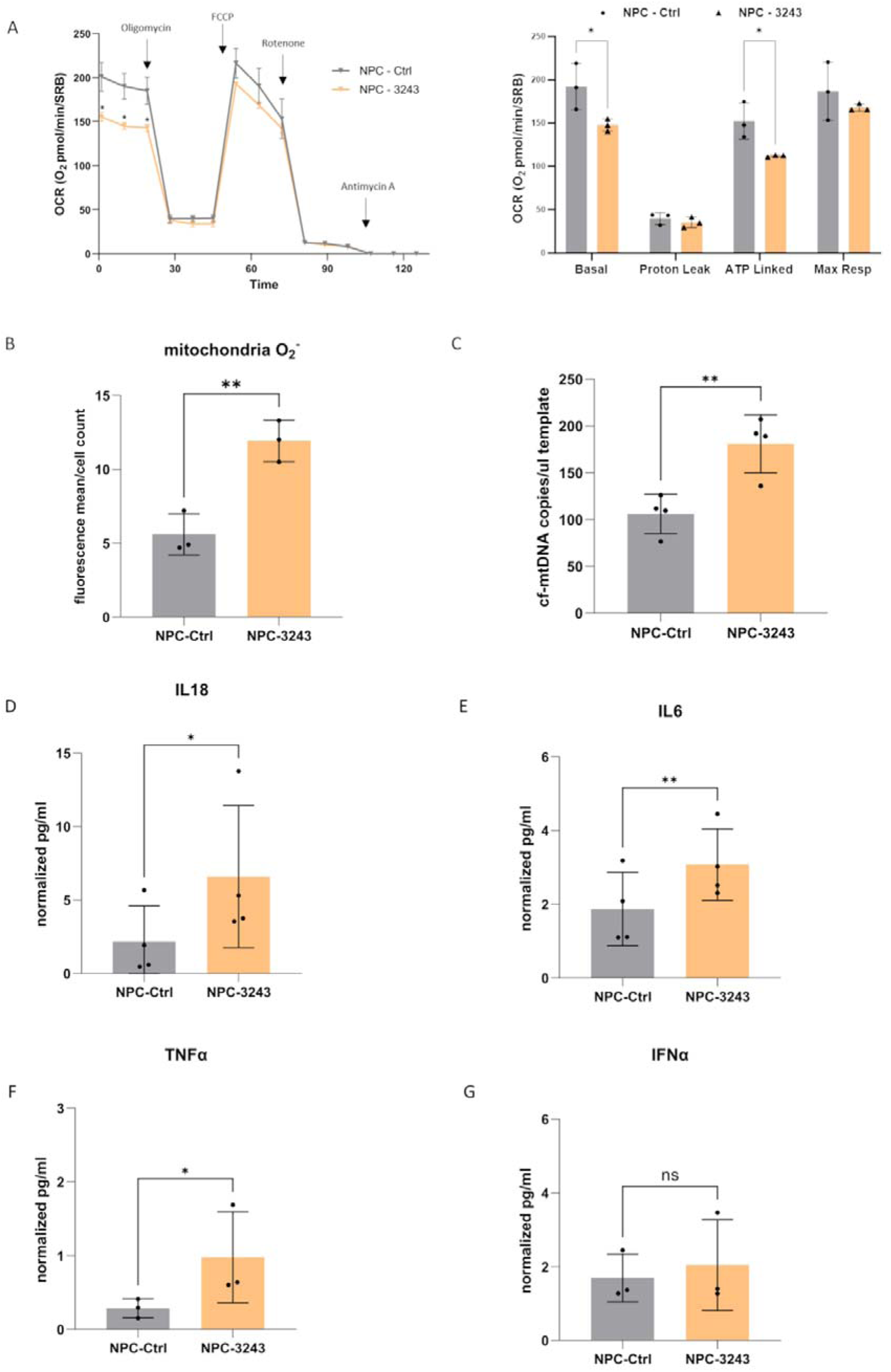
Characterization of the pathological phenotype in MELAS NPCs compared to the isogenic control.(A) Graphical representation of mitochondrial respiration, OCR was expressed as picomoles O2/min, normalized for protein content, under basal conditions and after injection of OXPHOS modulators (oligomycin, carbonyl cyanide 4-(trifluoromethoxy) phenylhydrazone (FCCP), rotenone, and antimycin A) in MELAS NPC (NPC-3243) versus isogenic controls (NPC-Ctrl). Besides, basal, proton leak, ATP-linked, maximal and spare respiration. All values are means and SD of 3 independent experiments (biological replicates). A two-way ANOVA with Šidák’s correction was used to analyze of the data. *, corresponding to an adjusted p-value of <0.03. **(B)** Quantification of mitochondrial superoxide by flow cytometry, based on MitoSOX™ mean fluorescence intensity normalized to cell count, from at least four independent biological replicates in MELAS NPC compared to isogenic control. An unpaired t-test was used for data analysis. ** corresponding to an adjusted p-value of <0.005 **(C)** The plot shows the cf-mtDNA level between MELAS NPC (NPC-3243) and isogenic control (NPC-Ctrl) for 4 independent biological replicates after 48h of growth in standard conditions. The data were normalized for the cells’ protein content. Data were analyzed by an unpaired t-test, ** indicates a p-value of 0.007. **(D-G)** Cytokine levels (IL-18, IL-6, TNFα and IFNα) were quantified in NPC-3243 and isogenic control cells. Data represent at least three independent biological replicates. Data were analyzed based on their distribution using a paired t-test or a ratio t-test. *, ** corresponding of adjusted p-value of <0.04 and of <0.001, respectively.

In summary, our results demonstrate a pronounced mitochondrial dysfunction in MELAS NPCs, characterized by impaired mitochondrial respiration and elevated mitochondrial oxidative stress. These metabolic alterations likely induce mitochondrial stress, promoting mtDNA release and subsequent activation of immune pathways and inflammation, even in NPCs, a cellular model closely resembling the disease-relevant target tissue.

### mtDNA release is strongly dependent on the presence of the m.3243A>G variant, elevated mitochondrial Ca^2+^ levels, and VDAC oligomerization

We studied a mutant cybrid cell line (3243-C) harboring 99% mutant heteroplasmy, taking advantage of the greater ease of manipulating these cells compared with primary fibroblasts or NPCs This analysis revealed increased cf-mtDNA levels under both basal and metabolically stressed conditions compared with *wild-type* cells (Fig. 7 A).

**Fig. 7.**
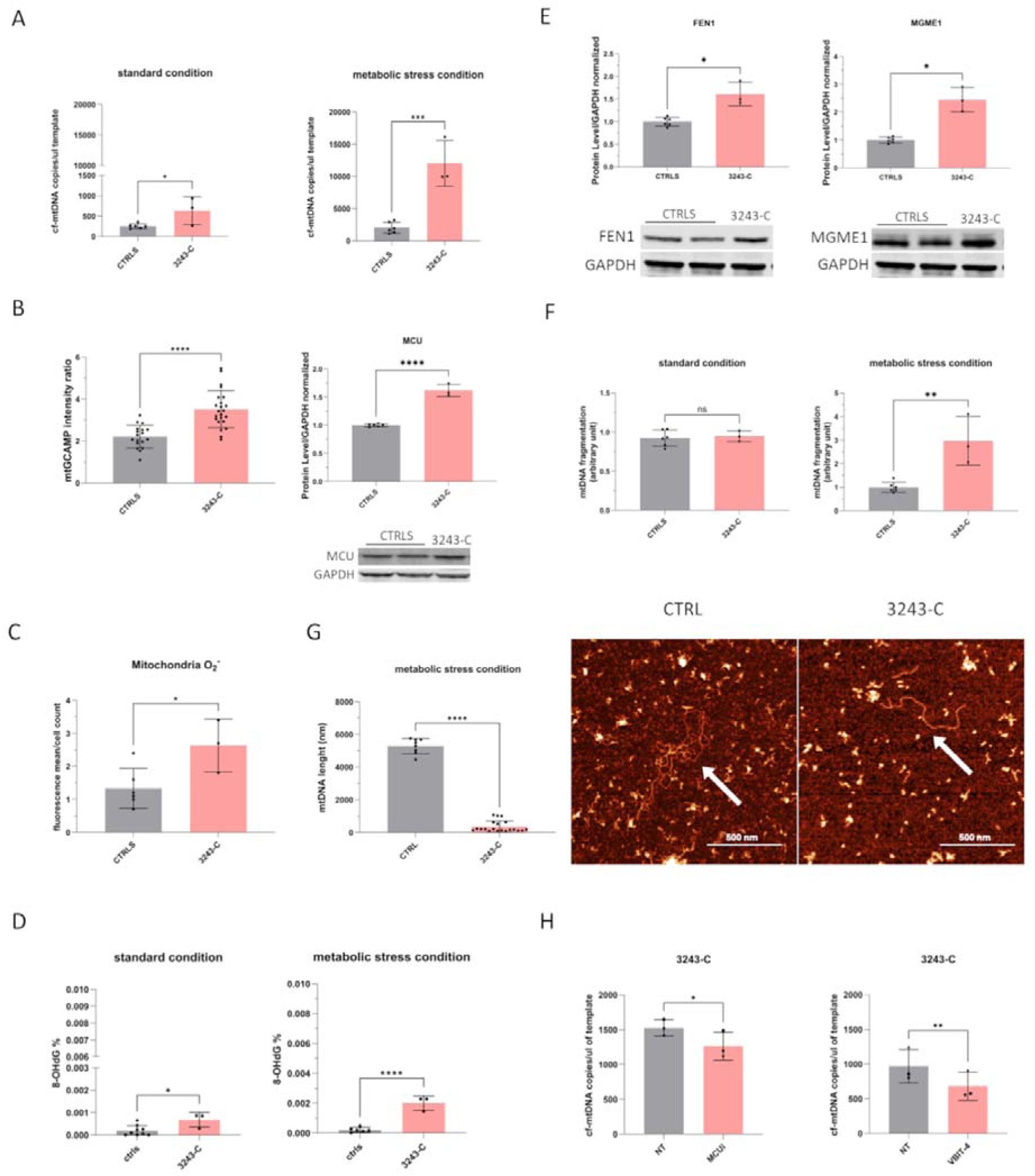
Evaluation of cf-mtDNA release and mitochondrial dysfunction in a cybrid cell model.(A) Assessment of cf-mtDNA levels in the culture medium after 16 hours of growth under standard and stress conditions. Data from three biological replicates per cell line are shown. **(B)** On the left, mitochondrial Ca^2+^ levels were assessed by fluorescence microscopy following transfection of cells with mtCGaMP. On the right, immunoblot analysis was performed to evaluate MCU protein levels. Representative images and densitometric quantification from at least three independent biological replicates are shown. GAPDH levels were used as a loading control for normalization. **(C)** Quantification of mitochondrial superoxide by flow cytometry, based on MitoSOX™ mean fluorescence intensity normalized to cell number, from at least three independent biological replicates. **(D)** Percentage levels of 8-OHdG in DNA extracted from mitochondria under standard and stress conditions in cybrids. **(E)** Immunoblot analysis was performed to assess FEN1 and MGME1 protein levels. Representative images and densitometric quantification from at least three independent biological replicates are shown. GAPDH levels were used for normalization. **(F)** Analysis of mtDNA fragmentation by capillary electrophoresis. The Y-axis shows the normalized ratio of the small peak area to the large peak area. Results are based on at least three biological replicates per cell line. **(G)** Histogram showing the distribution of mtDNA fragment lengths (nm) quantified by AFM, along with representative images illustrating mtDNA fragmentation under metabolic stress conditions in control and 3243-C cell lines, highlighting differences in mtDNA integrity. Reduced cf-mtDNA release in 3243-C cells compared to untreated cells after 16 h of treatment with MCUi-11 (2 μM) and VBIT-4 (10 μM). Statistical analysis for all data in this figure was performed using an unpaired t-test. *, **, *** and **** indicate p-values of <0.05, <0.001, <0.0002, and <0.0001, respectively.

Also in this model, the m.3243A>G variant enhanced mitochondrial Ca^2+^ uptake, as reflected by elevated mitochondrial Ca^2+^ levels and increased MCU protein abundance (Fig. 7 B), both significantly higher than in controls. Moreover, 3243-C cells displayed a mitochondrial stress phenotype characterized by increased mitochondrial superoxide production (Fig. 7 C).

A recently described mechanism proposes that oxidized mtDNA is cleaved by the endonuclease FEN1 into fragments of approximately 500–650 bp and released into the intermembrane space through the mPTP, whose opening is triggered by increased mitochondrial Ca^2+^ influx, and subsequently into the cytosol through a pore formed by VDAC oligomerization at the OMM ^51^. In the cybrid model, we therefore assessed mtDNA oxidation by isolating mitochondrial DNA and quantifying the percentage of 8-hydroxy-2’-deoxyguanosine (8-OHdG), a major oxidative DNA byproduct. Under both basal and metabolically stressed conditions, the percentage of 8-OHdG was significantly higher in the mutant line than in controls (Fig. 7 D).

Consistently, FEN1 protein levels were markedly increased in mutant cells, together with mitochondrial genome maintenance exonuclease 1 (MGME1), a key mitochondrial exonuclease involved in single-stranded DNA processing, which was also significantly upregulated in 3243-C cells, supporting enhanced mtDNA fragmentation (Fig. 7 E). Accordingly, DNA fragment analysis demonstrated increased mtDNA fragmentation in 3243-C cells under metabolic stress compared with controls (Fig. 7 F). These findings were further supported by atomic force microscopy (AFM), which showed increased mtDNA fragmentation in the MELAS cell line, in contrast to the predominantly full-length mitochondrial genome in control cells (Fig. 7 G).

To further validate the mechanisms underlying mtDNA release, we inhibited mitochondrial Ca^2+^ influx and VDAC oligomerization by treating cybrid cells for 16 h with either a MCU-complex inhibitor (MCUi11) ^54^ or a VDAC oligomerization inhibitor (VBIT-4) ^51^ at concentrations of 2 µM and 10 µM, respectively. Quantification of cf-mtDNA in the culture medium showed a significant reduction under both conditions (Fig. 7 H), indicating that mtDNA release relies on mitochondrial Ca^2+^ uptake and VDAC oligomerization.

## Discussion

This study highlights a novel molecular mechanism in cell models carrying the m.3243A>G pathogenic variant, consisting of enhanced mtDNA leakage from mitochondria into cytoplasm, and, ultimately, into the extracellular space. This is prompted by increased oxidative stress and consequent mtDNA oxidation and fragmentation, which, in turn, drive MCU-mediated mitochondrial Ca^2+^ accumulation and mtDNA release through VDAC. Although this process has been described for other pathological conditions ^55,56^, it represent an unappreciated mechanism in MELAS, potentially involved in its pathogenesis. In fact, the downstream cascade triggered by cf-mtDNA in the m.3243A>G model includes activation of the NLRP3 inflammasome and pyroptosis, leading to the secretion of pro-inflammatory cytokines (IL-1β and IL-18), thus suggesting a contributing role of inflammation in the progression of MELAS and non-MELAS phenotypes.

Consistent with our previous observation of increased ccf-mtDNA in affected patients ^19^, mutant fibroblasts exhibited elevated cf-mtDNA levels in the culture medium compared with controls, under both basal and stress conditions, as forcing the use of OXPHOS with galactose-medium. Notably, under this stress condition, both cell lines showed higher mutant heteroplasmy in cf-mtDNA than in intracellular mtDNA. However, this difference may be related to increase of dead cells with more dysfunctional mitochondria (i.e., higher mutant mtDNA heteroplasmy), as also indicated by the viability assay performed in galactose-containing medium. This process could account also for the observation of a lower intracellular m.3243A>G load in galactose-medium as compared with the standard growth condition. Thus, it is likely that this change in heteroplasmy may be due, at least partially, to this artifact. On the contrary, the difference in mutant heteroplasmy between intracellular and cf-mtDNA observed in the F#1 line under standard growth conditions (in absence of cell death) suggests that damaged mitochondrial genomes may be preferentially extruded from dysfunctional mitochondria. Variability in the distribution of the m.3243A>G variant within individual cells might mask this process, possibly explaining the different result observed in the F#2 line.

Immunofluorescence analysis further revealed increased cytoplasmic localization of mtDNA in mutant cells, indicating enhanced mtDNA release from mitochondria. Given that both cytosolic and extracellular mtDNA act as a DAMP ^57^, these findings suggest that mtDNA release may contribute to inflammatory signaling in MELAS. As expected, mutant fibroblasts displayed reduced viability under metabolic stress, impaired oxidative phosphorylation, compensatory mitochondrial biogenesis ^58^, and decreased mitochondrial membrane potential ^40^, consistent with persistent mitochondrial dysfunction ^41^. Altogether, these observations support a model in which dysfunctional mitochondria preferentially eliminate damaged mtDNA as a protective response, however, leading to the downstream activation of inflammatory pathways.

Transcriptomic profiling in fibroblasts revealed upregulation of cell death pathways, including pyroptosis, alongside a broad pro-inflammatory signature consistent with sterile immune activation. Key pyroptotic regulators (e.g., *IFI27, NLRP10* ^44,59^) and multiple cytokines (*IL-1*β*, IL-18, IL-6, TNF*α) were upregulated, whereas negative regulators of cytokine release, such as *SOCS2,* were downregulated ^46^. Pathway analysis further indicated activation of mitochondrial Ca^2+^ influx and outer mitochondrial membrane permeabilization, alongside downregulation of metabolic pathways, exacerbating mitochondrial stress.

Functionally, mutant fibroblasts exhibited Caspase-1 activation and increased secretion of IL-1β and IL-18, confirming NLRP3 inflammasome engagement ^16^. In parallel, increased IFNα, TNFα, and IL-6 levels, together with activation of RIG-I/MAVS and STAT1 signaling, indicate that mitochondrial nucleic acids also trigger interferon-mediated responses ^47^. These findings suggest that both mtDNA and mitochondrial RNA contribute to a multifaceted activation of innate immunity ^47^, particularly in the cell line from the patient suffering stroke-like episodes in analogy to other mitochondrial conditions ^60^.

Mechanistically, we identify mitochondrial oxidative stress and Ca^2+^ dysregulation as key drivers of mtDNA release. Mutant fibroblasts exhibited elevated reactive oxygen species and impaired antioxidant defenses, characterized by reduced GPX1 and catalase levels, both at mRNA and protein levels, suggesting a transcriptional rewiring potentially prompted by the m.3243A>G-dependent dysfunction ^58^. This imbalance likely promotes hydrogen peroxide accumulation and mtDNA oxidative damage. Consistently, we observed increased mitochondrial Ca^2+^ levels, together with upregulation of MCU and ITPR3, supporting enhanced Ca^2+^ transfer at mitochondria-associated membranes ^51^.

We next extended our investigations to MELAS patient-derived NPCs, a disease-relevant cell model, which recapitulated the fibroblasts’ mitochondrial phenotype, including impaired oxidative phosphorylation, increased mitochondrial superoxide production, and elevated cf-mtDNA release. The direct comparison with isogenic *wild-type* NPCs demonstrated that this mitochondrial phenotype, including the mtDNA release, is causally linked to the presence of m.3243A>G pathogenic variant. Moreover, the increase in IL-18, IL-6, and TNFα suggests activation of innate immune and inflammasome pathways, potentially triggered by mitochondrial dysfunction and mtDNA release.

To further establish the prevalent role of mutant mtDNA over the nuclear genome, we employed a mutant cybrid cell model (3243-C, 99% heteroplasmy), which confirmed that the m.3243A>G variant alone is sufficient to induce mtDNA release from mitochondria with a different nuclear genetic background. In fact, these cells exhibited increased mitochondrial Ca^2+^ uptake and elevated oxidative stress, accompanied by higher levels of mtDNA oxidation, as indicated by increased 8-OHdG level. In line with recent reports ^51,61^, we observed upregulation of the endonucleases FEN1 and MGME1 and increased mtDNA fragmentation within mitochondria under metabolic stress, supporting a mechanism in which oxidized mtDNA is processed into smaller fragments prior to release. The lack of detectable fragmentation in standard conditions may reflect rapid extrusion of limited amounts of fragmented mtDNA, whereas under stress conditions, fragmentation exceeds the extrusion capacity and accumulates within mitochondria.

Pharmacological inhibition of MCU (MCUi-11) or VDAC oligomerization (VBIT-4) significantly reduced cf-mtDNA release, demonstrating that mtDNA extrusion directly depends on mitochondrial Ca^2+^ overload and outer membrane permeabilization. Together, these findings support a model in which oxidative stress-induced mtDNA damage promotes Ca^2+^-dependent mPTP opening and VDAC-mediated release of fragmented mtDNA. However, the lack of complete reversal of mtDNA extrusion may have different explanations. First, the complete pharmacological inhibition of MCU and VDAC oligomerization possibly needs higher concentrations, which, however, have cytotoxic effects, at least in the cybrids model. On the other hand, additional channels may be involved in the mtDNA release in MELAS cell models.

Translating our *in vitro* findings to the *in vivo* scenario, we propose that this mechanism may contribute to tissue-specific and systemic inflammation in patients carrying the m.3243A>G pathogenic variant. We hypothesize that during SLEs, exacerbated mitochondrial stress and pyroptotic activation may contribute to local tissue damage and neuronal loss. In patients not experiencing SLEs but suffering a more chronic slow deterioration of the CNS, ultimately leading to dementia, this neuro-inflammatory mechanism may be equally active as documented in our patient who did not suffer SLEs (F#1). Furthermore, as we previously documented ^19,20^, the high levels of circulating cf-mtDNA in plasma may have long-range effects by triggering a cytokine storm, which can involve peripheral organs and worsen clinical outcomes, culminating in multi-organ failure. This is supported by recent studies showing type I interferon signatures and increased inflammatory signaling in patients with mitochondrial diseases ^62^, as well as by evidence from animal models, including *Ndufs4 /* (Leigh syndrome ^63^) and *POLG1* mutated mice ^64^, in which inflammation contributes to disease progression. Notably, anti-inflammatory and immunosuppressive treatments, including corticosteroids, have shown clinical benefit in subsets of patients with mitochondrial disorders, including MELAS ^65–67^. More broadly, chronic inflammation, particularly neuroinflammation driven by activated microglia and astrocytes, is increasingly recognized as a key contributor to neurodegeneration across multiple disorders, including Alzheimer’s disease, Parkinson’s disease, amyotrophic lateral sclerosis, and multiple sclerosis ^68^. Our study provides solid evidence for druggable pathways warranting further investigation, with novel therapeutic approaches potentially shaping the natural history of MELAS.

While our findings provide consistent mechanistic evidence across complementary cell models, limitations include the reliance on *in vitro* systems and a limited number of patient-derived lines, which may not fully capture the complexity and heterogeneity of MELAS *in vivo*.

In conclusion, our findings define an oxidative stress– Ca^2+^–mtDNA release axis that directly links mitochondrial dysfunction to innate immune activation, suggesting that pharmacological targeting of mtDNA release downstream inflammatory pathways may represent a promising strategy to modify the disease course in MELAS and related disorders.

## Data availability

Data are available in the main text and in the supplementary materials. Additional datasets are available on Zenodo: https://doi.org/10.5281/zenodo.20412724.

## Supporting information

Supplementary results Figs. S1 to S6

## Acknowledgments

We sincerely thank all the patients for their invaluable participation and collaboration, as well as the Italian patient organization for mitochondrial diseases, Mitocon Onlus, for its continuous support. We are also grateful to Andrea Martinuzzi and Monica Montopoli (University of Padova) for kindly providing the control cybrid cell lines.

## Author contributions

Conceptualization: AM, MM

Methodology: PP,AM, MM, MT, AS, LC, MC, GC

Investigation: MM, AR, CF, DO, CVT, FV, AD, SP, CLM, VC, APP

Visualization: MM, AR Supervision: AM

Writing—original draft: MM, AM, VC, AR, SP, AD, FV, CVT

Writing—review & editing: all authors

## Supplementary Materials

See Supplementary Materials file.

