## Supplementary results Figs. S1 to S6 for "Cell-free mtDNA drives pyroptosis and inflammation in m.3243A>G mitochondrial disease models"

Moresco Monica *et al.*

**This file includes:**

Supplementary results

Figs. S1 to S6

Supplementary Text

**Results**

**Clinical History of patient F#1**

This proband is a 68-year-old male carrying the m.3243A>G/tRNA^Leu^ point mutation, with a heteroplasmy of 20% in blood cells and 48% in skeletal muscle.

The family history was relevant for the recurrence of diabetes, sensor-neural deafness, and migraine along the maternal line. The patient suffered migraine without aura since he was 18 years old. Deafness was evident at 20 years of age, subsequently requiring a prosthesis at 64 years. Diabetes started at 20 years of age, requiring insulin since the onset. Since the age of 50, he has also suffered fatigue and proximal hyposthenia, which worsened over time. At 58 years of age, he presented 4 episodes of chest pain that were interpreted as Non-ST-Elevation Myocardial Infarction, and a diagnosis of Ischemic Cardiomyopathy was made at that time. He also presented progressively worsening renal insufficiency since he was 56 years old (stage IV when he was 67).

A progressive worsening of cognitive functions was also evident since the age of 64, associated with aggressive behaviors, and at the last evaluation (at 67 years of age), the Mini-Mental State Examination (MMSE) was 18/30.

He also presented at 68 years (March 2024) with an acute rhabdomyolysis with associated worsening of renal function, paralytic ileus, and pneumonia. Neurological functions also deteriorated on that occasion with episodes of hallucinations and psychomotor agitation. Electroencephalogram (EEG) and Computed Tomography (CT) scan were negative for stroke-like episodes. Blood exams showed persistent lactic acidosis in 2022, requiring IV correction with bicarbonate. Brain Magnetic Resonance Imaging (MRI) with spectroscopy (MRS) performed in October 2023 (at 67 years, supplementary figure 1A) showed diffuse cortical atrophy, including also hippocampus and putamen, cerebellar atrophy, and signs of vascular leukoencephalopathy. Diffuse basal ganglia (caudate and pallidal) calcifications were also evident. MRS showed a pathological lactate peak. Respiratory function tests showed a moderate-to-severe restrictive defect, not requiring ventilatory support. Electromyography showed diffuse myopathic signs.

He died in August 2024 due to status epilepticus in the absence of stroke-like episodes at brain MRI complicated by acute ingested pneumonia and worsening of renal function.

**Clinical History of patient F#2**

The patient presented with psychomotor delay and moderate intellectual disability. He also suffered from an autism spectrum disorder since childhood. Woll-Parkinson-White was diagnosed when he was 3.

Since he was 5 years old, he started presenting migraine attacks. Since adolescence, he has presented with hypoacusia. Hypertrophic cardiomyopathy was diagnosed when he was 20 years old.

The first left parieto-temporal stroke-like episode (SLE) occurred in 2010 (at 23 years of age). At that time, MELAS was suspected and confirmed by genetic testing, which showed the presence of the m.3243A>G/tRNA^Leu^ pathologic variant. Muscle biopsy showed the presence of 2 Cytochrome c oxidase (COX) negative fibers and 1 Ragged Red Fibers (RRF). Other SLEs occurred in 2014 at 26 years (left Parietal Temporal Occipital, PTO) and 2 episodes in 2015 at 27 years (right frontal and right PTO, see supplementary figure 2A-B). Investigations performed at 27 years showed:

-increased lactate levels (28 mg/dl, nv 5-22 mg/dl)

-MRI (see images): right PTO SL lesion

-EEG: epileptiform discharges in the right posterior regions

Diabetes was diagnosed when he was 34 years of age. In 2020, in the context of COVID-19 infection, he presented respiratory and heart failure with the necessity of tracheostomy and intensive care unit hospitalization. The month after he presented a severe worsening of the heart function, requiring an implantable cardioverter defibrillator. Amiodarone therapy was added. Renal insufficiency also occurred.

The patient died at 36 years of age on Oct 13^th^, 2023, due to severe heart failure with subsequent multi-organ failure in the course of a severe thyrotoxicosis secondary to amiodarone.

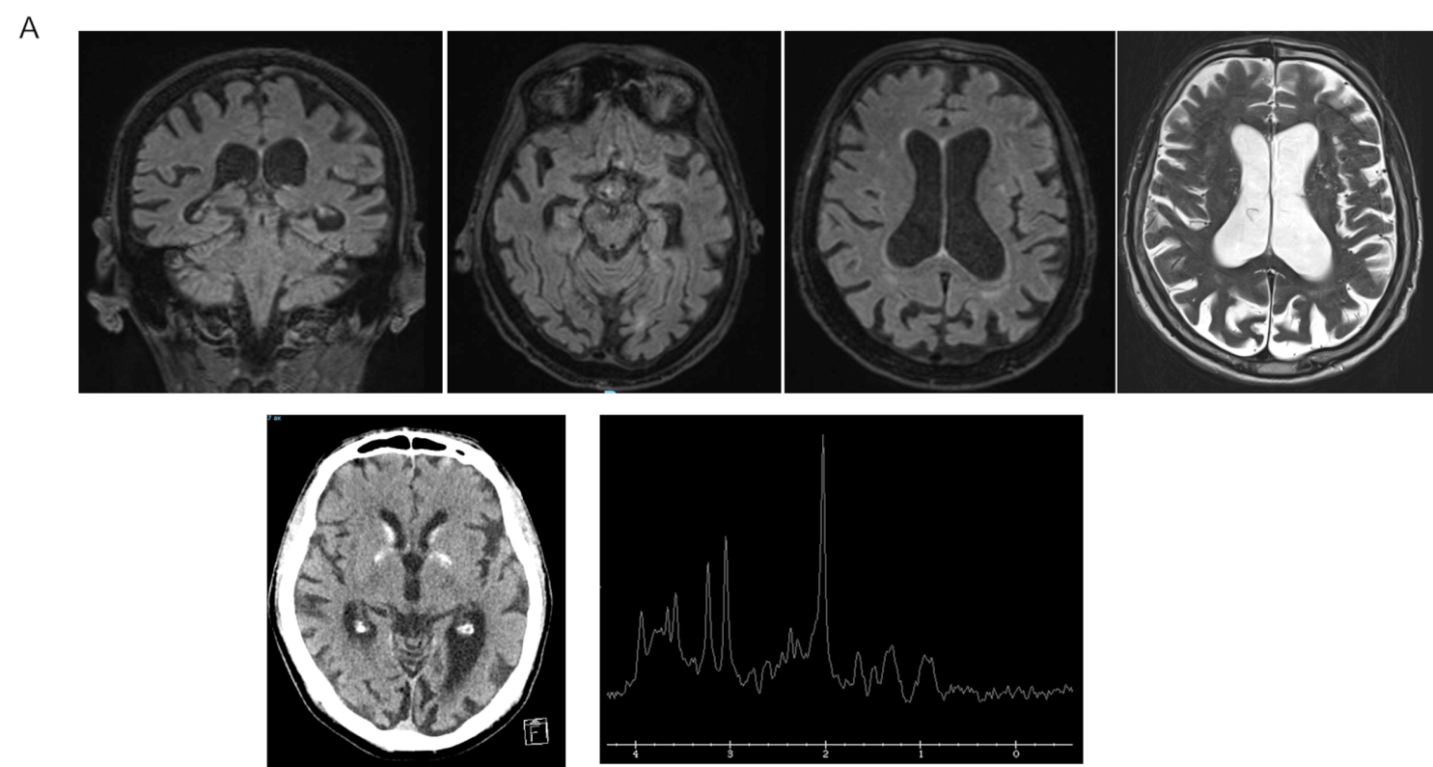

Fig. S1.

**(A)** Brain MRI with spectroscopy (MRS) performed in October 2023 on patient F#1.

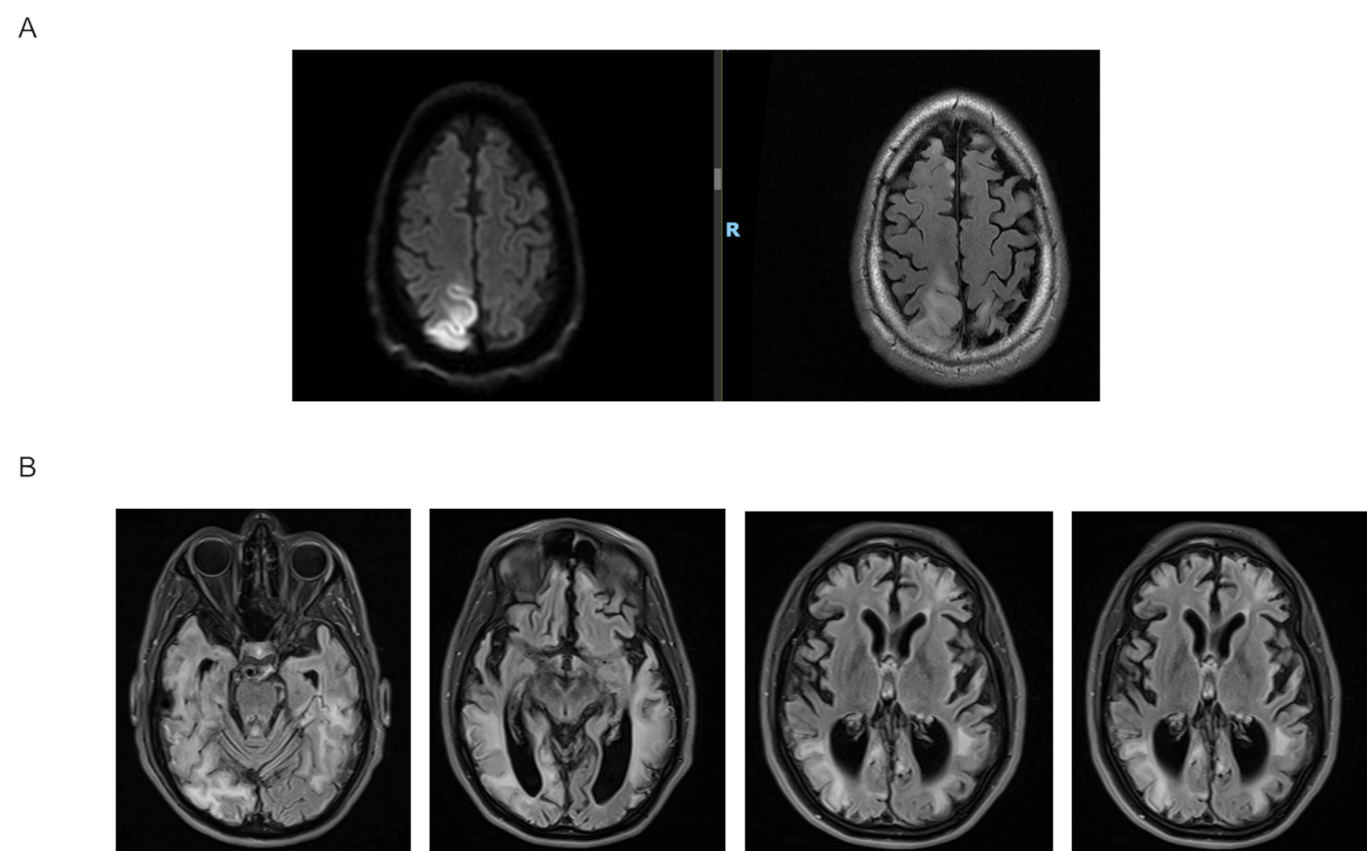

Fig. S2.

**(A)** Brain MRI with spectroscopy (MRS) performed in April 2015 with Diffusion-Weighted Imaging (DWI) and Fluid-Attenuated Inversion Recovery (FLAIR) on patient F#2.

**(B)** Last brain MRI on 2017 of patient F#2.

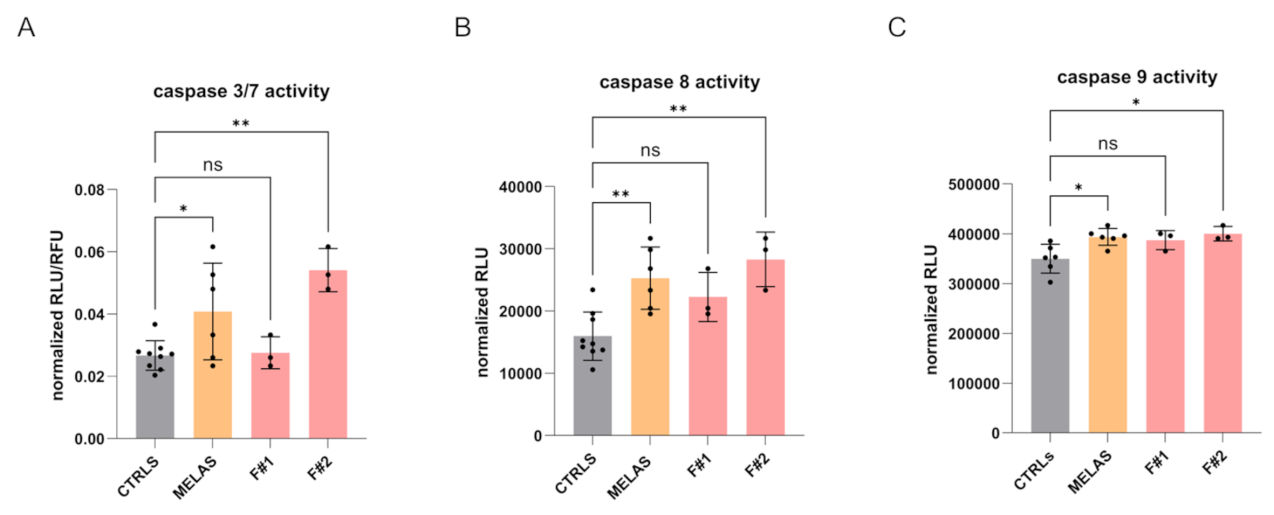

Fig. S3.

**(A)** Evaluation of caspase-3/7 activity in fibroblast cell lines. Values shown in the graph represent the relative light units (RLU) normalized to relative fluorescence units (RFU).

**(B)** Evaluation of caspase-8 activity in fibroblast cell lines. The results shown in the graph represent the RLU mean ± SD of three biological replicates for each cell line normalized on protein content.

**(C)** Evaluation of caspase-9 activity in fibroblast cell lines.

The results shown in the graph represent the RLU mean ± SD of three biological replicates for each cell line normalized on protein content. All analyses were performed using one-way ANOVA followed by Dunnett’s multiple-comparison test.

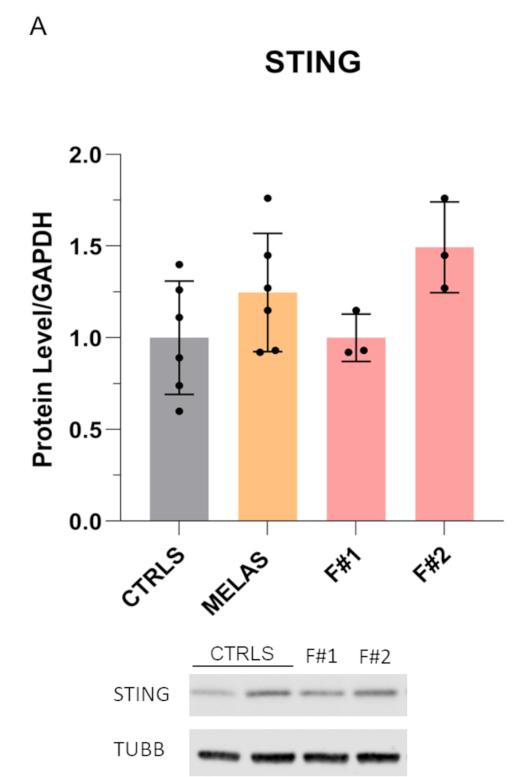

**Fig. S4.**

**(A)** Immunoblot analysis for the evaluation of STING protein level. Representative image and densitometric quantification from at least three independent biological replicates are shown. TUBB levels were used as a loading control for normalization.

For all data presented in this figure, statistical analysis was performed using one-way ANOVA followed by Dunnett’s multiple-comparison test.

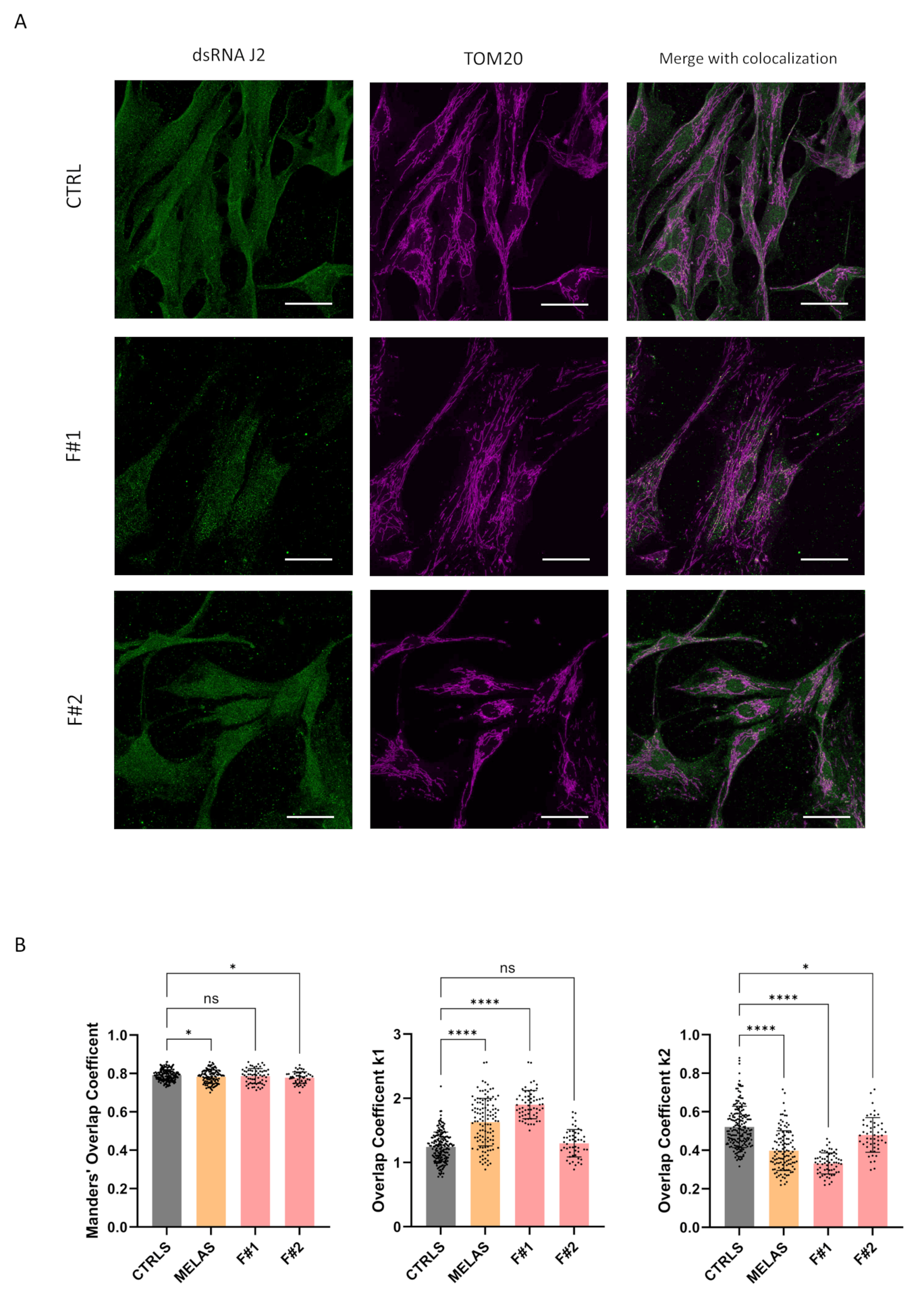

**Fig. S5.** **Evaluation of dsRNA J2 colocalization in mitochondria on MELAS fibroblasts.**

(A) Representative confocal images showing co-localization in fibroblasts under standard conditions, stained with an anti-dsRNA J2 antibody (green) and anti-TOM20 antibody (pink) to label mitochondria. Scale bar: 40 μm.
(B) Overlap coefficient according to Manders (MOC), and overlap coefficients k1 and k2 between MELAS fibroblasts and matched controls following 48h of growth in standard condition. Data were analyzed by one-way ANOVA test corrected for multiple comparisons (Dunnett) for MOC coefficient, whereas for k1 and k2 coefficients we used Kruskall-Wallis test corrected for multiple comparisons (Dunn’s), *, **, and **** indicate significant differences from the corresponding controls, p-value of 0.04, 0.001 and <0,0001, respectively.

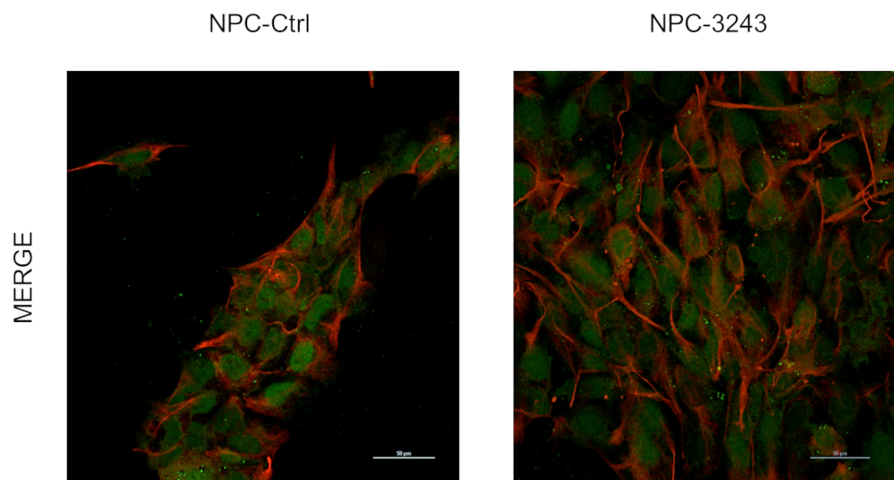

**Fig. S6.** NPC characterization in mutant (NPC-3243) and isogenic *wild type* cells (NPC-Ctrl) via Immunofluorescence. Sox2 (green) and Nestin (red) were used respectively as self-renewal, and immature neuroepitelial cell marker.
